# The interplay between growth rate and nutrient quality defines gene expression capacity

**DOI:** 10.1101/2021.04.02.438188

**Authors:** Juhyun Kim, Alexander P.S. Darlington, Declan G. Bates, Jose I. Jimenez

**Affiliations:** Department of Chemical Engineering, Imperial College London, South Kensington Campus, London, SW7 2AZ, United Kingdom; School of Engineering, University of Warwick, Coventry, CV4 7AL, United Kingdom; Department of Life Sciences, Imperial College London, South Kensington Campus, London, SW7 2AZ, United Kingdom

**Keywords:** ribosomes, cellular economics, metabolic constraints, (p)ppGpp

## Abstract

The gene expression capacity of bacterial cells depends on the interplay between growth and the availability of the transcriptional and translational machinery. Growth rate is widely accepted as the global physiological parameter controlling the allocation of cell resources. This allocation has an impact on the ability of the cell to produce both host and heterologous proteins required for synthetic circuits and pathways. Understanding the relationship between growth and resources is key for the efficient design of artificial genetic constructs, however, it is obscured by the mutual dependence of growth and gene expression on each other. In this work, we investigate the individual contributions of molecular factors, growth rate and metabolism to gene expression by investigating the behaviour of bacterial cells growing in chemostats in growth-limited conditions. We develop a model of the whole cell that captures trade-offs in gene expression arising from the individual contributions of different factors, and validate it by analysing gene couplings which emerge from competition for the gene expression machinery. Our results show that while growth rate and molecular factors, such as the number of rRNA operons, set the abundance of transcriptional and translational machinery available, it is metabolism that governs the usage of those resources by tuning elongation rates. We show that synthetic gene expression capacity can be maximised by using low growth in a high-quality medium. These findings provide valuable insights into fundamental trade-offs in microbial physiology that will inform future strain and bioprocesses optimisation.

## Introduction

The ultimate goal of synthetic biology is to engineer reliable and robust genetic circuits. These circuits have a range of potential applications in biomedicine, industry and environmental science^1^, but ensuring their reliable performance requires precise control over gene expression. To this end, it is crucial to understand the inherent constraints on gene expression created by the limited cellular economy, in order to allow optimal allocation of resources between host and circuit genes. Recently, researchers have observed that the expression of a particular gene can affect the activity of another seemingly unconnected gene due to sharing of the cell’s resources. Such resource-coupled gene expression is mainly caused by limitations in the numbers of ribosomes^2-8^. Several approaches have been developed to reduce this coupling^4,6,8,9^, for instance, we successfully mitigated competition for ribosomes in co-expressed genes using quasi-orthogonal ribosomes combined with a resource allocation controller^8^.

The association between cellular growth and ribosome abundance has previously been investigated in order to understand how cells allocate their protein synthesis capacity. As the synthesis of ribosomes is energetically costly, cells need to maintain an optimal concentration of these macromolecules to maximise cellular fitness^10,11^. In bacteria, evolutionary trade-offs have shaped the number of rRNA operons to achieve optimal ribosomal biosynthesis. For instance, wild type *Escherichia coli* carry seven copies of the operons and adapt to nutritional perturbations more quickly that other isogenic strains with different copy numbers of the rRNA operons^12^. The cellular concentration of ribosomes is also controlled by the nutritional stress-induced alarmone (p)ppGpp^13-17^. As a response to amino acid starvation (the stringent response), cells accumulate (p)ppGpp which represses the expression of both rRNAs and ribosomal-protein coding genes through direct interaction with the RNA polymerase^18^.

Bacterial growth rate acts as a global regulator of ribosomal concentrations by coordinating the distribution of the cellular proteome. This allocation has been described by partitioning the *E. coli* proteome into three fractions: the R fraction (ribosomes/ribosome-associated), the E fraction (containing metabolic enzymes), and a growth independent fraction^19-21^. Cells allocate more of their proteome to ribosome-associated fractions under nutrient rich conditions to support maximum growth, and thus lower amounts of metabolic proteins are produced. In contrast, under poor nutrient conditions, cells synthesize more metabolic proteins to produce ATP at the expense of ribosome production, which leads to slower growth^19-21^. Consequently, the synthesis of ribosome-associated proteins competes with the production of non-ribosomal proteins.

To investigate such cellular fitness strategies, growth rate (and hence ribosome biosynthesis) has been manipulated by altering nutrient quality^22-24^. However, this can potentially produce confounding results, due to the impact of different metabolic effects on the resource allocation. For example, different carbon sources could lead to distinct profiles of amino acid synthesis which in turn could affect translation beyond the availability of ribosomes. In addition, the impact of growth rate on such resource-mediated coupling has so far not been explored, and this question is crucial to ensuring that multiple genes are expressed at the right combination and concentration under dynamic resource allocation regimes.

In this work, we investigate resource allocation in the cellular proteome by using a continuous culturing system that enables a systematic investigation of different growth scenarios for the same metabolic conditions. By combining theory and experiments, we obtain novel insights into the interplay between transcription and translation and the role of rRNA operons. These insights allow us to untangle the relationship between growth and metabolism to identify the impact on gene couplings of both carbon sources and the (p)ppGpp network. These results shed new light on fundamental trade-offs in microbial physiology and will enable optimised cultivation of microbial cell factories for diverse biotechnological applications.

## Results

### A dynamic model of metabolism, resource allocation and cellular growth

To investigate the above questions, we developed a simplified model of the whole cell that accounts for multiple trade-offs in gene expression. By combining mechanistic models of gene expression and natural feedback with phenomenological models of growth and a simple representation of metabolism we produced a tractable model that is capable of recapitulating the different distributions of cellular proteins. This model goes beyond existing ones^9,25^ by accounting for the following key universal constraints: (i) finite biosynthetic capacity, (ii) competition for total number of proteins, (iii) competition for RNA polymerases by mRNA and rRNA promoters, (iv) competition for ribosomes by mRNAs, (v) resource biosynthesis (both production of RNA polymerase and ribosomes, including their autocatalysis) and (vi) dynamic growth-based feedback.

A schematic of the full model is shown in Fig. 1 and full mathematical details are described in the Supplementary Material. Here we summarise the main features and species within the model. The model considers the growth of cells in the presence of a single external substrate (*s*_*x*_). This substrate is imported into the cell by transporter proteins (*p*_*T*_) to produce *s*_*i*_, which itself is converted by enzymes (*p*_*E*_) into an expanded metabolism consisting of a proxy ‘energy’ species (*e*), amino acids (*a*) or nucleotides (*n*). The production of *a* and *n* are dependent upon both the internal substrate *s*_*i*_ and energy *e*. Gene expression dynamics are captured by a simple four state model for each gene. Firstly, the RNA polymerase (*P*) reversibly binds to free promoters to form transcription complexes (*K*_*x*_, where *x* is the gene of interest). These are ‘consumed’ by transcription *T*_*X*_(⋅) to produce messenger RNAs (*m*_*x*_). The mRNA are reversibly bound by ribosomes (*R*) to produce the translation complex (*c*_*x*_). The mRNA are released upon termination of translation at rate *T*_*L*_(⋅) which produces new proteins (*p*_*x*_). The transcription [translation] process is dependent upon both the energy species and nucleotides [amino acids]. We modeled gene expression regulation by scaling the RNA polymerase-promoter association rate as detailed in the Supplementary Material. All genes are regulated by the cell’s internal energy status. The dynamics of resource biosynthesis mirrors those of other genes. We modelled ribosome biosynthesis as a two-step process consisting of transcription of ribosomal RNAs and transcription/translation of ribosomal proteins. These bind to production functional ribosomes. The model was fit as described in the Methods to the resource economy data^26^ and successfully recapitulates the key associations between growth and resource production (Figure S1A) as well as predicting peptide elongations (Figure S1B)

**Figure 1.**
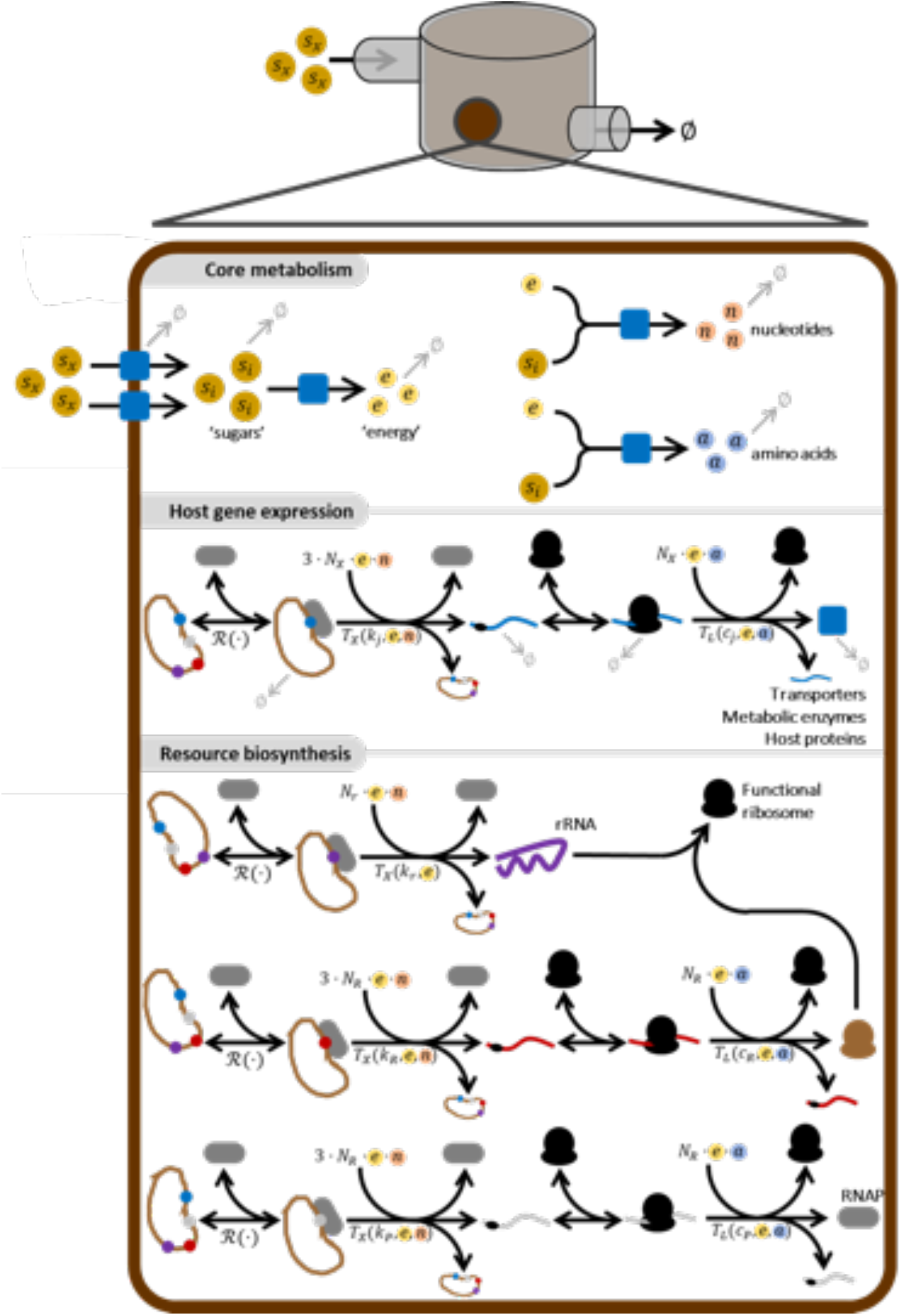
Model schematic. The ordinary differential equation model describes a simple cell with a four-state metabolism, gene expression and resource biosynthesis. The external nutrient (*s*_*x*_) is imported by transporter proteins (*p*_*r*_) to become the internal substrate (*s*_*i*_). This is converted to an energy source (*e*) by enzymes (*p*_*E*_). These enzymes also convert the internal metabolite to nucleotides (*n*) and amino acids (*a*) in reactions which also utilise the energy substrate. All host genes (such as transporters, enzymes and other host proteins) are expressed in a three-step process: each gene (*j*) is bound by an RNA polymerase to form a transcription complex (*K*_*j*_) which undergoes transcription (*T*_*X*_), consuming energy and nucleotides in the process, to produce an mRNA (*m*_*j*_). The mRNA is then bound by a ribosome to form translation complex (*c*_*j*_) which undergoes translation (*T*_*L*_), consuming energy and amino acids in the process, to produce a protein (*p*_*j*_). The model explicitly considers resource biosynthesis. The RNA polymerase is produced like other host proteins. Ribosomes are produced in a three-step process. First, ribosomal proteins are produced in the same manner as other host proteins. Secondly, rRNAs are produced by transcription utilising RNA polymerase, energy and nucleotides in the same manner as other host RNAs are produced. The ribosomal proteins then bind the rRNA to form functional ribosomes which are capable of translation. *N*.*B*. All reactions are described in detailed in the Supplementary Material. Not all reactions are shown to simplify the schematic.

### Growth rates set gene expression trade-offs in carbon-rich conditions

It is well established that changing nutrients within a growth media impacts both cell growth rate and ribosome content, with both being correlated^27,22,28^. Moreover, the interplay between gene expression and growth on a range of media has previously been established^22-24^. However, in these experiments the changes to the cell’s metabolism (such as changing carbon source) cause concurrent changes to growth rate, resource biosynthesis and gene expression. To dissect these relationships independently we utilized a chemostat to allow for the tight control of bacterial growth in glucose rich conditions. This allows the manipulation of bacterial growth rates and the quantification of gene expression levels under identical external nutrient conditions. To investigate resource-mediated gene coupling we utilized an *E. coli* strain that carries a plasmid encoding a constitutively expressed GFP and an inducible RFP under the control of LuxR (activated by acyl-homoserin lactone; AHL). This circuit enables a quantitative assessment of gene coupling and resource utilization as described previously^9^. Cells were cultured at the following growth rates: 0.1, 0.3, and 0.5 h^−1^, as determined by the dilution rate of the minichemostat and in batch culture, using a range of AHL concentrations in each of those conditions. After five generations of growth, resulting in a physiologically steady state of the culture^29^, the production of the reporters at each given growth rate was measured by using flow cytometry.

When the chemostat was set at a 0.1 dilution rate, cells in the culture displayed a higher synthetic gene expression activity, compared to those in cultures with higher growth rates (Fig. 2A). The expression levels of GFP and RFP were three- and four-fold higher, respectively, in cells with a 0.1 h^−1^ growth rate, compared to those in the culture set at a 0.5 h^−1^. Cells grown in glucose-limited batch culture showed the highest growth rates but the lowest activity of the two reporters (Fig. 2A). We measured ribosome production by labelling the ribosomal L9 subunit with monomeric superfolder GFP (msfGFP) and found increased fluorescence with increased growth rate, reflecting a higher ribosomal protein synthesis (Fig. 2B). Therefore, the reduced synthesis of the reporters at high growth rates is not due to a lower production of ribosomes. These results are in agreement with previous observations^23^ and are supported by our resource allocation model (Fig. 2C), which shows that at higher growth rates cells invest more in cellular resources, with RNA polymerase and ribosomes preferentially producing their own components, and a concurrent decrease in circuit mRNA and protein synthesis (Figs. 2D and S2).

**Figure 2.**
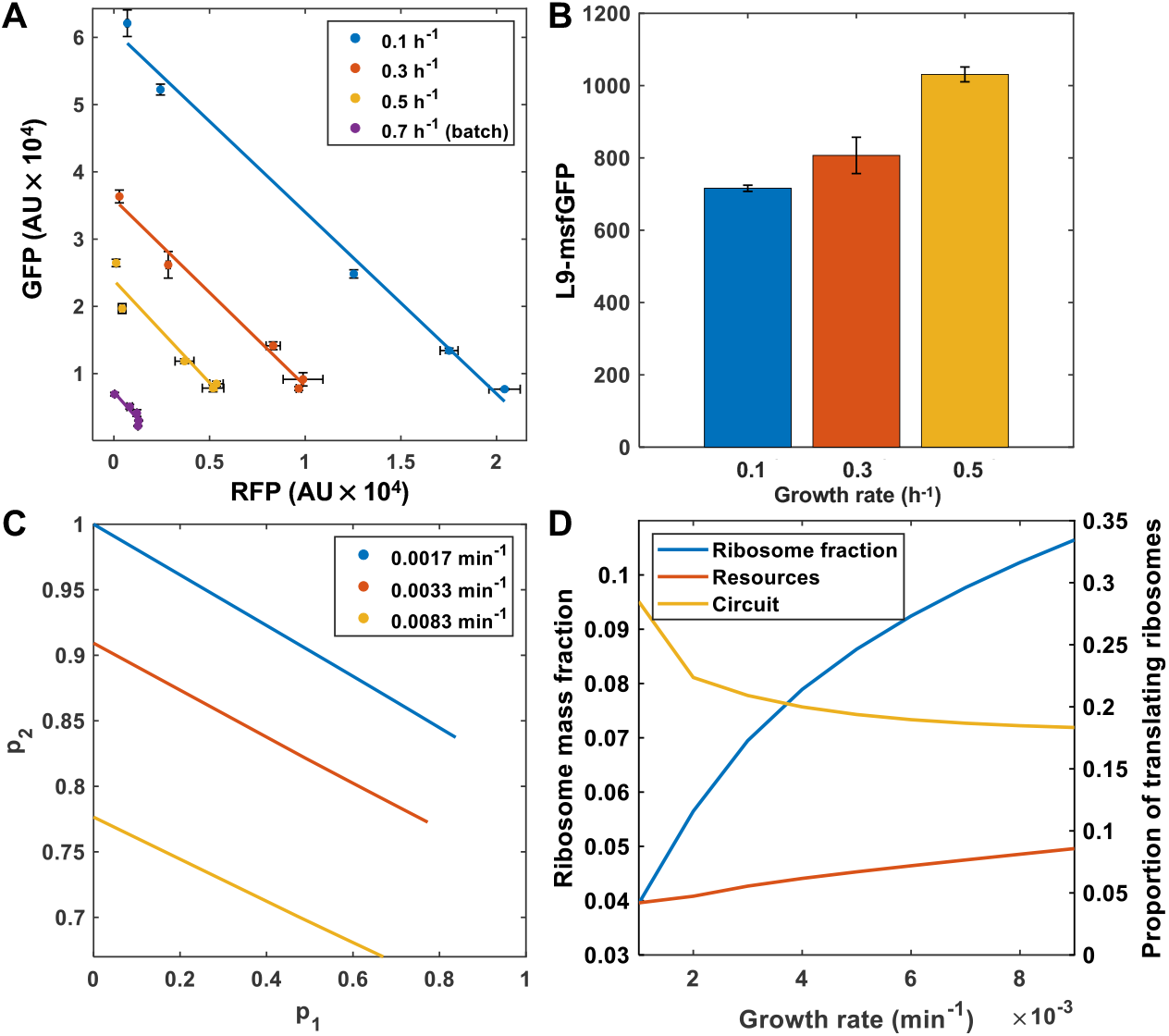
Growth-rate dependence of gene expression and resource allocation. **(A)** Resource mediated gene expression couplings were examined for different growth rates and AHL concentrations in cells carrying a genetic circuit that allows co-expression of GFP and RFP, which were determined by flow cytometry. Cells were also grown in batch cultures to quantify the reporter expression at the maximum growth rate. **(B)** Ribosomal protein production was quantified by monitoring the expression, for different growth rates, of a genomic fusion of the ribosomal protein L9 to msfGFP. **(C)** Simulations of the steady state concentration of the two gene circuit, normalized by maximum protein production, for different growth rates. Isocost lines were simulated by maintaining varying *u*_1_ = 0 to 1 and maintaining *u*_2_ = 1. Simulations were carried out as described in the Methods. **(D)** Simulations of the cell’s internal resource economy expressing two circuit genes (with induction constants of *u*_1_ = 0.1, *u*_2_ = 1) over a range of growth rates.

Our experimental system also enabled us to test whether the growth rate affects gene expression in resource-mediated competition. As shown in Fig. 2A, where the gene expression coupling occurred in continuous cultures; the expression level of GFP decreased when RFP accumulated in the cells regardless of the growth rate, and the concentration of the reporters is constrained by a linear manifold known as an ‘isocost line’^5^. The same level of coupling, defined as the ratio of the trade-off in the expression between the two genes, was maintained in all the conditions tested (Fig. 2A) suggesting that the cost of the protein production (as determined by the slope of the linear manifold – a measure of the resources required per protein produced^5^) is independent of the growth rate and is governed only by the abundance of ribosomes.

### Effect of selective inhibition of transcription or translation on resource allocation

Next, we sought to investigate the selective inhibition of components of the gene expression machinery to understand the effects of resource allocation at defined growth rates. To this end, sublethal concentrations of either rifampicin (Rif) or chloramphenicol (Cm) were added to the continuous culturing system for partial inhibition of, respectively, RNA polymerases or ribosomes. Again, the use of chemostats allows us to untangle the impact of resource inhibition whilst forcing cells to maintain a fixed growth rate (which is not possible in a batch culture system).

When cells were treated with Rif, lower intensities of both GFP and RFP were recorded, with expression levels that gradually decreased with increasing concentrations of the antibiotic (Fig. 3A). We modelled the inhibition of RNA polymerase by sequestering RNA polymerase through varying *K*_*rf*_ within the model (Fig. 3B; see Supplementary Material for details). As *K*_*rf*_ is increased, the number of RNA polymerases and ribosomes dedicated to their own production increases (Fig. 3C). This results in a concurrent fall in production of other genes; i.e. the proportion of RNA polymerases [ribosomes] transcribing [translating] circuit genes falls (Fig. S3). This increase is required to maintain growth at the set rate (Fig. 3C). We confirmed this prediction by treating cells grown at 0.1 h^−1^ carrying the L9-msfGFP fusion with Rif. This demonstrates increased synthesis of ribosomal proteins with increasing condition of Rif addition (Fig. 3D). Similarly, the production of both reporters decreased when a sublethal concentration of Cm was added to the culture (Fig. S4). Our model demonstrates that a simple natural feedback emerging from resource autocatalysis (i.e. that resources are required to make themselves) is sufficient to explain the fall in circuit expression which occurs during resource-inhibition (Figs. S5 and S6). We also found that partial inhibition of both transcription or translation at a fixed growth rate did not cause a significant change in coupling (i.e. the isocost gradients are constant), rather, there was only a change in the amount of circuit protein produced, indicating that reducing the transcription or translation activities does not change the cellular resource allocation regime.

**Figure 3.**
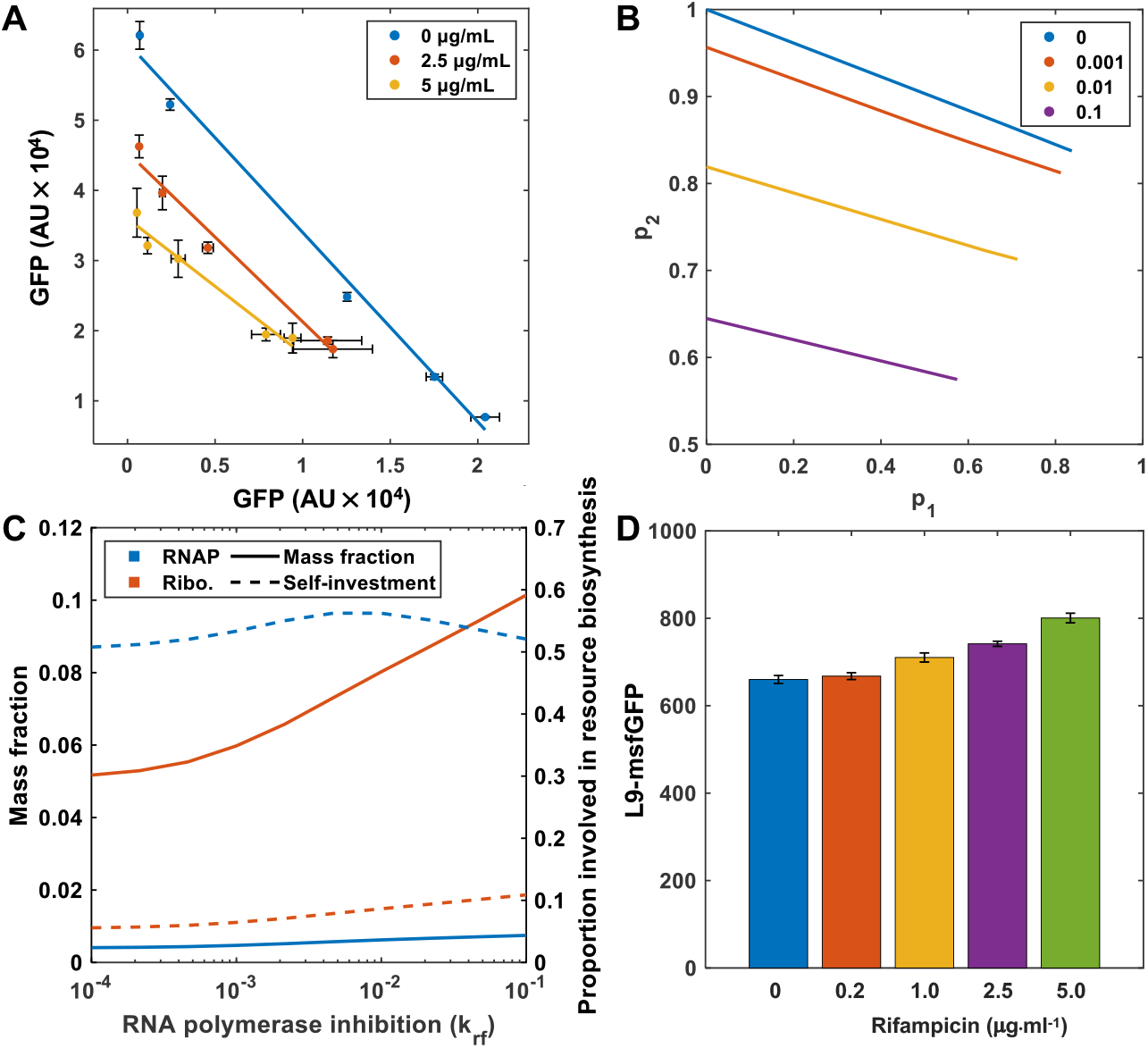
Effect of the selective inhibition of transcription on resource availability. **(A)** The strain carrying the GFP and RFP reporters was cultured in the minichemostat in the presence of sublethal concentrations of rifampicin and at different growth rates. **(B)** Simulations of the steady state concentration of the two reporter circuit normalized by maximum protein production, for different antibiotic transcriptional inhibition constant ranging from *K*_*rf*_ 0 to 0.1. Isocost lines were simulated by maintaining varying *u*_1_ = 0 to 1 and maintaining *u*_2_ = 1. Simulations were carried out as described in the Methods section. **(C)** Simulations of the cell’s internal resource economy expressing the two-reporter circuit (with induction constants of *u*_1_ = 0.1, *u*_2_ = 1) over a range of RNA polymerase inhibition constant *K*_*rf*_ varied on a log10 scale from −4 to −1. **(D)** The synthesis of the L9 ribosomal protein was examined in different concentrations of rifampicin at a fixed growth rate.

### The interplay between rRNA transcription, growth and gene expression

We investigated the effect of tuning the maximum size of the ribosomal pool on the protein production budget by using strains with successive deletions of the *rrn* operons^30^. Strains with different numbers of *rrn* operons were transformed with the resource-sensitive RFP-GFP circuit plasmid to measure the spare translation capacity and cultivated in the chemostat at either 0.1 or 0.5 h^−1^ growth rates, representing slow and fast growth conditions, respectively. When the production of GFP was analysed, a higher accumulation of the protein was observed in the cells growing with the low dilution rate compared to the high dilution rate, regardless of their *rrn* copy number (Fig. 4A) – demonstrating again that growth and circuit gene expression are inversely correlated. The analysis also showed that the expression levels of GFP in fast-growing cells were weakly correlated with rRNA gene dosage. In contrast, the synthesis of GFP was strongly dependent on the copy number of the *rrn* operon in slow-growing cells (Fig. 4A). The synthesis of the fluorescent protein dropped by 30% and 60% in the strains containing one and two deleted *rrn* operons, respectively, compared to the wildtype strain growing at the same growth rate of 0.1 h^−1^. Nonetheless, only the two initial *rrn* deletions affected gene expression significantly, whereas the impact of further deletions was negligible (Fig. 4A). Our model predicts that this fall in expression is due to an increased self-investment effect; as the number of rRNA genes falls, rRNA transcription falls and so the number of functional ribosomes falls. Therefore, to maintain the desired growth rate, each functional ribosome spends more time translating ribosomal proteins. As the number of rRNA genes is scaled down from 100% to 0% of its original literature value the ribosomal mass fraction increases (Fig. 4B). This effect is predicted to be more pronounced at higher growth rates.

**Figure 4.**
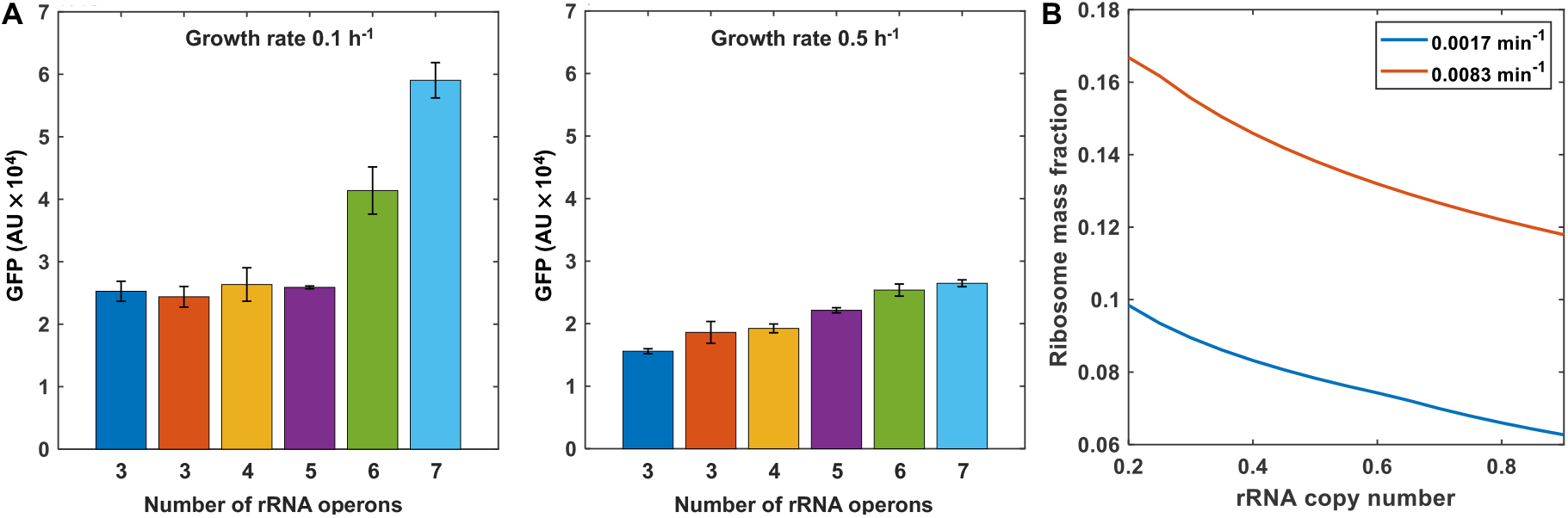
Translational capacity dependent upon rRNA operon copy numbers. **(A)** Constitutively expressed GFP levels were measured in strains carrying different number of *rrn* operons by growing them in the continuous culture at either low growth rate (0.1 h^−1^; left panel) or high growth rate (0.5 h^−1^; right panel). The strains containing ribosomal deletions used in this experiment were (from left to right) SQ78, SQ52, SQ49, SQ40, SQ37 from ^30^ and the wildtype. **(B)** Simulations of the cell’s ribosomal mass fraction of cells expressing the one gene circuit where the rRNA copy number (*g*_*r*,0_) is scaled between 0 and 1 of its wild-type value. Simulations were carried out as described in the Material and Methods section.

### Metabolism plays a key role in the allocation of cellular resources

Previous work has investigated nutrient limitations as one of the key factors controlling ribosome synthesis^31,32^, although in these batch growth studies the contribution of nutrients is not isolated from the effect of changes in growth rate. To dissect the distinct contributions of growth rate and metabolism on gene expression, we grew the RFP-GFP reporter carrying strain at a set growth rate with different nutrient conditions. Samples were taken and circuit protein expression assessed by flow cytometry. At constant growth rate in glucose cultures, we found that increasing the nutrient richness by supplementing with casamino acids did not affect the expression of circuit genes (Fig. S7). However, reducing nutrient quality by cultivating cells in glycerol, instead of glucose, led to a 30% reduction in the reporter activity, compared to that in the cells using glucose as sole carbon source (Fig. 5A). To identify the mechanistic cause of this fall in circuit expression we modelled the impact of changing nutrient quality by varying *ϕ*_*e*_ (which governs the *s*_*i*_ to *e* conversion rate), *ϕ*_*a*_ (which governs the *s*_*i*_ and *e* to *a* conversion rate) and *ϕ*_*n*_ (which governs the *s*_*i*_ and *e* to *n* conversion rate) both independently and in combination. We found that varying *ϕ*_*a*_ alone is sufficient to recapitulate the experimental result, suggesting that the impact of reducing the carbon source from glucose to glycerol reduces the availability of translational substrates, such as amino acids (Fig. 5B). Assessing the model’s internal cellular economy and elongation rates showed that when *ϕ*_*a*_ falls there is a concurrent fall in elongation rate, which is both energy and amino acid dependent (Fig. 5C). To maintain growth rate, the mass fraction of ribosomes rises due to increased resource investment in resource production (Fig. 5C, Fig. S8E and F). To corroborate this prediction, we measured the production of the L9 ribosomal protein and, as expected, glycerol-grown cells exhibited higher levels of fluorescent intensity relative to those in glucose-grown cells (Fig 5D). This result suggests that the lower expression of circuit genes observed in the cells grown in glycerol is due to a subtle interaction between elongation rate and number of ribosomes, and that growth rate does not set resource levels *per se*. Rather, growth rate sets the global translation rate, which is the product of the elongation rate and the number of translating ribosomes (Eq. 25 in the Supplementary Materials). As the elongation rate falls (e.g. due to the poorer quality substrate, the number of translating ribosomes must rise for growth to be maintained.

**Figure 5.**
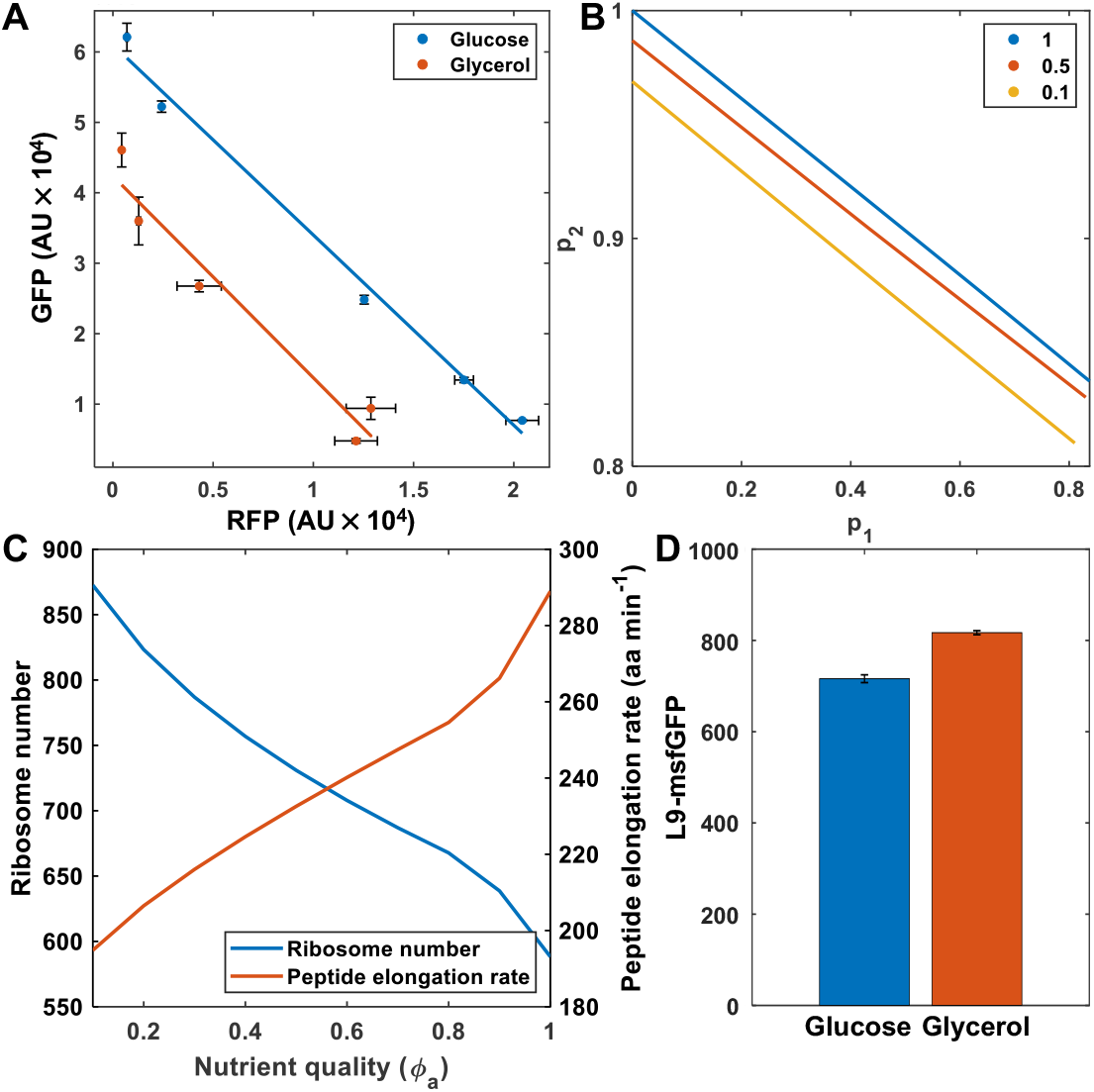
Nutrient quality effect on the allocation of cellular resources. **(A)** The expression capacity of the circuit genes was analysed by comparing the data of the fluorescent intensity obtained from the dual fluorescent reporter strain grown either in glucose (Figure 1A) or in glycerol as the carbon source. **(B)** Simulations of the steady state concentration of the two-reporter circuit, normalized by maximum protein production, for different nutrient quality. Different nutrient qualities were simulated by scaling *ϕ*_*a*_ form 0.1 to 1 of its nominal value. Isocost lines were simulated by maintaining varying *u*_1_ = 0 to 1 and maintaining *u*_2_ = 1. Simulations were carried out as described in the Material and Methods section. **(C)** Simulations of the cell’s internal resource economy and peptide elongation rate of a cell expressing the two gene circuit (with induction constants of *u*_1_ = 0.1, *u*_2_ = 1). **(D)** The expression level of the L9 ribosomal protein was measured in the same culture condition as described in (A).

### (p)ppGpp affects the global distribution of the proteome

Our findings show that the allocation of ribosomes to expendable proteins is inversely correlated with the growth rate, in agreement with the growth laws reported previously^19-21,23,24^. A salient question is what are the effects of physiological mechanisms that are known to play a role in the allocation of cellular resources on this trade-off. Ribosome synthesis is mainly controlled by the alarmone, (p)ppGpp^15,17,18^. (p)ppGpp accumulates at high levels when nutrients are limited, which results in the repression of both ribosomal proteins and rRNAs, and a decrease in growth rates^15,17,18^. In *E. coli*, (p)ppGpp is synthesised by RelA and degraded by the hydrolase SpoT^14,18^. We studied the role of the alarmone on resource allocation for set growth rates, monitoring the activity of the RFP-GFP circuit in either *relA* or *spoT* deleted strains, exhibiting low or high (p)ppGpp levels, respectively^33,34^. When the mutant strains, carrying the reporter genes, were cultured in the chemostat at a growth rate of 0.1 h^−1^, a two-fold decrease in fluorescent intensity was recorded for both GFP and RFP in the *relA* deleted strain compared to its counterpart *spoT* mutant strain (Fig. 6). This result suggests that the accumulation of the alarmone due to the mutation in the degradation activity encoded by *spoT* results in more resources being available for the circuit genes, due to a lower investment in ribosome synthesis. Interestingly, the RelA deficient strain showed a reduced resource-mediated coupling, which is associated with changes in the ratio between mRNAs and ribosome abundance (Fig. 6)^5^. This indicates that the alarmone is not only involved in controlling ribosome concentration, but also in global transcriptional suppression^17,35^ and thus the mutant strain exhibits a different cost of protein synthesis^34^. To identify the cellular processes that could have the largest effect on the gradient of the isocost line, we carried out a global sensitivity analysis of key metabolic parameters within the model^36^. We found that both the gradient and y-intercept of the isocost line are most sensitive to the *o*_*X*_ parameter (Figure 6B and C, respectively). This parameter represents the transcriptional energy threshold of the enzymes – i.e. the level of the internal energy species at which the transporter and enzyme production regulation is half maximal. We varied this parameter by +/−50% to assess the impact on the isocost line and found that decreasing *o*_*X*_ (i.e. making the enzymes more sensitive to the cell’s internal energy supply) results in a decrease in the circuit output (Figure S9). We also find the isocost line is also sensitive to the energy threshold of the RNA polymerase (*o*_*P*_). Our global sensitivity analysis also identifies the elongation rate thresholds (*k*_*X*_, *k*_*r*_ and *k*_*γ*_) as key determinants of the isocost line features. This suggests that the effect of the *relA* and *spoT* mutations on the isocost line is through a global reallocation of transcription and potential changes in the elongation process.

**Figure 6.**
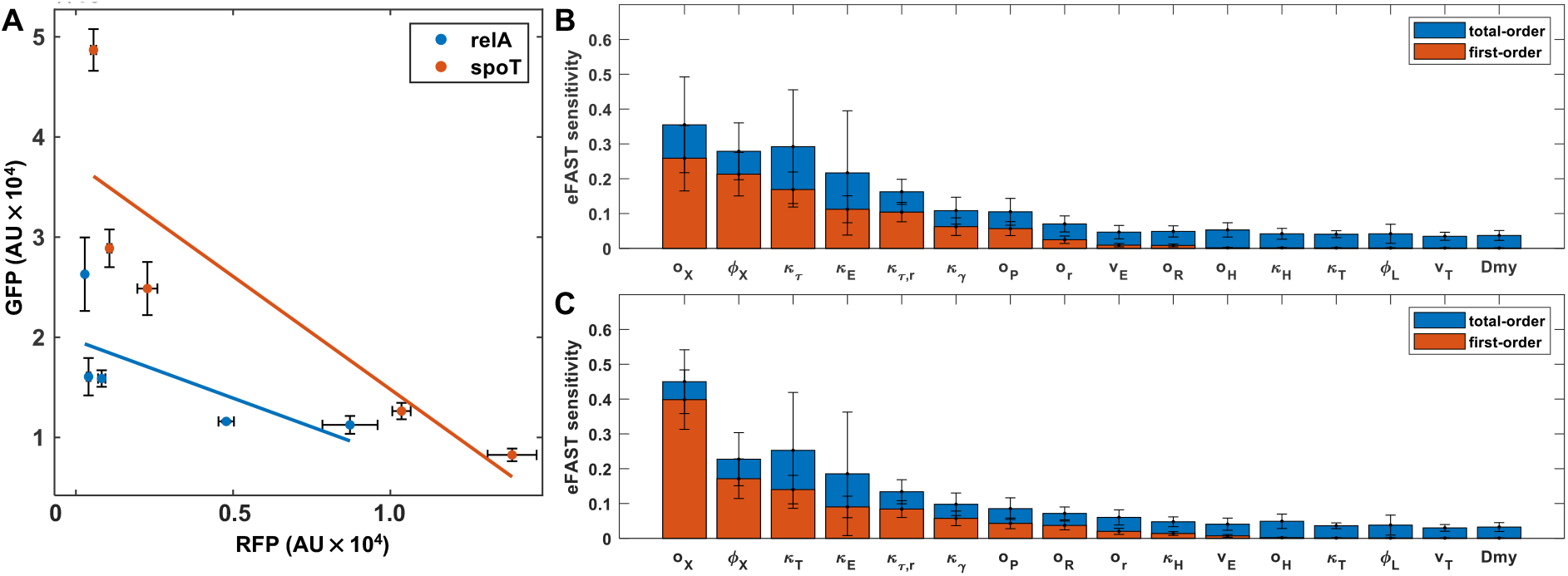
(p)ppGpp mediated allocation of resources for gene expression. **(A)** Resource mediated gene expression coupling was examined in mutants of the (p)ppGpp alarmone network. A *relA* mutant produces lower levels of (p)ppGpp than the wildtype strain while the *spoT* mutant accumulates more (p)ppGpp. Strains were cultured at a fixed growth rate in the minichemostat. **(B)** The global sensitivity of the isocost line gradient to each parameter was calculated using the eFAST method^36^ using a parameter perturbation of up to 10%. **(C)** The global sensitivity of the isocost line y-intercept to each parameter was calculated as described in the main text.

## Discussion

In this study, we investigated how cellular resources are allocated to circuit gene expression under different cellular economic regimes. In artificial systems where ribosomal synthesis is not coupled to growth – e.g. cell-free extracts or cells endowed with orthogonal ribosomes – translational capacity for circuit genes is directly proportional to the size of the ribosomal pool. However, in natural systems, bacterial ribosomes preferentially synthesize more ribosomes to obtain rapid growth. This is at the expense of the production of other proteins, including those of synthetic circuits or pathways. This strategy is defined by bacterial growth laws which address the partitioning of the cellular proteome, as first proposed by Hwa and colleagues^19-21^. In their original work, they cultivated *E. coli* cells in media containing varied carbon sources, resulting in different growth rates to obtain insights into the relation between the growth rate and the proteome distribution.

In this work we analysed the specific contributions of metabolism and growth to the allocation of resources for gene expression. To this end, we exploited a continuous culture system that allowed us to manipulate the growth rate while maintaining the carbon source, to enable the investigation of *both* growth-dependent and growth-independent effects on resource allocation. Under growth-limited (i.e. growth controlled) conditions, we found lower expression levels of circuit genes in fast-growing cells relative to those in slow-growing cells, which is in line with previous observations from carbon-limited conditions. Our model analysis showed that this is the result of a greater allocation of cellular resources to the synthesis of gene expression machineries, which is required for cells to achieve maximum growth. This strategy enables cells to maximise reproductive fitness. It is noteworthy that the cost of protein synthesis (i.e. the resources required per protein produced) was barely changed when cells redistributed resources, and thereby the level of resource mediated gene expression coupling was maintained.

This resource allocation strategy allows microbial cells to maintain cellular fitness in harsh conditions. For example, when transcription was partially inhibited with rifampicin, ribosomal protein synthesis increased, while the expression level of the circuit genes decreased. The model showed that this is the result of the self-investment of the gene expression machinery in its own synthesis to support the set growth rate. Similarly, we observed lower synthesis of the reporters in the presence of chloramphenicol-mediated translational inhibition. It is worth noting that although phenomenologically the cells behave in the same way, the specific regulatory mechanisms of resource (re-)allocation may vary depending on the point of inhibition. When treated with rifampicin, cells produce more ribosomes at the expense of the synthesis of other cellular RNAs, including circuit mRNAs, which leads to a reduced allocation of the proteome to heterologous genes. On the other hand, the inhibition of translation results in a reduced number of active ribosomes as evidenced in previous reports conducting polysome profiling^22,37^. Those studies show that when the cells are cultured in the presence of Cm, they accumulate 70S monosomes, which are not significantly involved in protein synthesis.

Our results show that this relationship between growth rate and resource allocation is also affected by other factors. We found that as the *rrn* operon copy number increased, circuit gene expression increases in both slow and fast growth cells, but this effect is more pronounced in slow growth (where cells are investing less in resource production). The alarmone (p)ppGpp also plays an important role by setting the number of resources available for the synthesis of circuit proteins. We determined that when cultured at the same growth rate, a *relA* deficient strain, containing low (p)ppGpp levels, synthesized fewer heterologous proteins compared to a *spoT* mutant. Impairing RelA activity leads to reduced (p)ppGpp production, potentially raising ribosome synthesis while impairing SpoT, leading to (p)ppGpp accumulation and inhibition of ribosome synthesis^14,15,18^. As ribosome synthesis falls, circuit gene expression rises, thus explaining the different behaviour of these mutants.

We dissected the different contributions of growth rate and carbon source to the allocation of the cellular budget for gene expression. We determined that gene expression is not exclusively tuned by the number of ribosomes, but also depends on the peptide elongation rate. We found that increasing the quality of an already rich medium through amino acid supplementation did not change the translational capacity of the cell to produce the reporter proteins. On the other hand, when the cells metabolized glycerol, a poor-quality substrate, a lower expression of the circuit genes was observed compared to that in glucose (a high-quality carbon source *for E. coli*). These results are not due to a reduced availability of ribosomes in glycerol, and on the contrary, ribosomal protein synthesis was increased in glycerol. Instead, our model suggests that this is the result of a decrease in the translation elongation rates in the poor nutrient condition. In poor quality nutrients, such as glycerol, cells produce fewer translational building blocks – such as tRNA, elongation factor Tu, and guanosine triphosphate (GTP)^22^. Our model predicts that to compensate for the lower elongation rate and to maintain the externally set growth rate, cells synthesize more ribosomes in order to maintain their total translational capacity. This results in a reduction in spare capacity and so there is a reduction in the synthesis of non-ribosomal proteins.

The impact of host circuit interactions due to finite cellular resources has been blamed on the failure of synthetic gene circuits since the field’s inception. Recently these interactions (notably in terms of ribosome-mediated coupling between co-expressed genes) have begun to be rigorously quantified. However, our results show that in addition to ribosomal competition, metabolism and (p)ppGpp are key drivers in setting the cell’s synthetic budget. These results have significant implications for the design of synthetic circuits and biotechnological pathways and imply that recombinant protein production can be increased by simple strategies such as identifying the carbon sources enabling the best elongation rates and artificially reducing the growth rates. The conceptual framework and results described here represent a step towards a complete understanding of the impact of this hitherto neglected layer of host-circuit interactions.

## Materials and methods

### Minichemostat set up

The custom-made mini continuous-culture system consisted of 15 reactors was set following the protocol^29^ with some modifications. We used 80 ml glass bottles (VWR, Radnor, PA, USA) for the 50 ml cultures. We used silicone tubing (ID, 4 mm; OD 6 mm), 90 degree plastic tubes (ID, 4 mm; OD 6 mm), Norprene pump tubes (ID, 1.6 mm; OD, 5 mm, Sigma-Aldrich, St. Louis, MO, USA), plastic connectors, luer fittings, and syringe needles (14G and 19G) for the inflow and outflow of liquid cultures through a multi-channel peristaltic pump (Cole-Parmer, St. Neots, UK). A magnetic bar was used in each reactor for continuous culture agitation with a microplate stirrer (2mag, Munich, Germany). The pump flow rate was set to 80, 230 and 400 µl·ml^−1^ to generate growth rates of, respectively, 0.1, 0.3 and 0.5 h^−1^.

### Cultivation of bacterial cells

Experiments in this study were conducted with *E. coli* MG1655^38^ and derivatives (Table S1). Strains were grown in batch or continuous cultures using M9 minimal medium (6 g·l^−1^ Na_2_HPO_4_, 3 g·l^−1^ KH_2_PO_4_, 1.4 g·l^−1^ (NH_4_)_2_SO_4_, 0.5 g·l^−1^ NaCl, 0.2 g·l^−1^ MgSO_4_·7H2O) supplemented with glucose (0.2%) and casamino acids (0.2%) at 30°C, unless otherwise indicated. The antibiotics kanamycin (50 µg·ml^−1^), ampicillin (150 µg·ml^−1^) and gentamicin (20 µg·ml^−1^) were added when necessary.

Chemostat cultures carrying the pSEVA63-Dual plasmid were initiated from overnight cultures in batch. Right after inoculation cultures were induced with N-acyl homoserine lactone (AHL, Sigma-Aldrich, St. Louis, MO, USA, final concentrations of 0.5, 1, 5, 10 nM) and/ or sublethal concentration of antibiotics such as rifampicin and chloramphenicol were added. After five generations of growth, when the population of cells in the chemostat reached a steady state, samples were collected and to determine fluorescent reporter expression levels (see below). *E. coli* MG1655 L9-*msf*GFP harbouring the L9 ribosomal protein labelled with GFP was cultured in the minichemostat without inducers.

### Cloning procedures and construction of reporter strains

The characteristics of the bacteria, plasmids and primers used in this study are described in Table S1. DNA manipulation was carried out following standard protocol^39^. Plasmid DNA was isolated from bacterial cells using a commercial QIAprep Spin Miniprep Kit (Qiagen, Hilden, Germany). Restriction endonucleases were purchased from New England Biolabs (NEB, Ipswich, MA, USA).

The *E. coli* MG1655-derived strains containing the *rplI(L9)-msfGFP* fusion was constructed with a seamless allelic replacement method described previously^40,41^. The delivery plasmid pEMG^41^ was used for the genomic exchange *rplI* x *rplI*-*msf*GFP and was built as follows: First, the entire *rplI* gene (TS1^*rplI*^, ∼0.5 kb) and downstream (TS2 ^*rplI*^, ∼0.5 kb) regions of the 3’-end of the gene were amplified with primer pairs rplI-TS1F/R and rplI-TS2F/R respectively (Table S1). Another DNA segment bearing the *msfGFP* sequence flanked by overlapping regions with those of the TS1^*rplI*^ and TS2^*rplI*^ fragments was prepared by amplification of plasmid pBG with primers rplI-msfGFP-F/R. The resulting TS1^*rplI*^, *msfGFP* and TS2^*rplI*^ were then joined by isothermal assembly^42^ and cloned in pEMG (Table S1) previously digested with *Eco*RI and *Bam*HI, to yield pEMG-*rplI*-*msfGFP*, transformed into *E. coli* DH5α λ*pir*. This plasmid was transferred to *E. coli* MG1655 where it can not replicate and was co-integrated in the chromosome. Next, the pSW plasmid that expresses I-SceI endonuclease under the *Pm* promoter^43^ was introduced by electroporation into the pEMG-*rplI*-*msf*GFP cointegrated. Selected clones were grown in LB medium with Ap and 3MBz (5 mM) to activate the *Pm* promoter, allowing I-SceI expression, which in turns forces a second recombination event by cleaving the co-integrate. Cells were plated on LB agar and the generation of a chromosomal *rplI*-*msf GFP* fusion was confirmed by testing the loss of the pEMG-*rplI*-*msf* GFP-encoded Km resistance gene. Km-sensitive clones were selected and chromosomal replacement further confirmed by PCR with oligonucleotides rplI-TS1F/TS2R (Supplementary Table S1). pSW was finally cured by serial dilutions in the absence of antibiotics, yielding the strain *E. coli* MG1655 *rplI-msf GFP*.

To inactivate the SpoT activity in the BW25113 strain, the target genomic region was replaced with a kanamycin antibiotic cassette (pKD4) using primers spoT KO F/−R, following a method described previously^44,45^ and the genomic deletions were further confirmed by PCR.

### Fluorescence measurements

Intracellular abundance of fluorescence proteins was determined with an Attune NxT Flow Cytometer (ThermoFisher, Waltham, MA, USA) analyzing GFP and RFP expression using blue (excitation 488 nm; emission 530/30 nm) and yellow (excitation 561 nm; emission 620/15 nm) lasers respectively. 100 µl of the sample was taken from each culture and mixed with 200 µl of PBS prior to injecting into the flow cytometer. 30,000 events of each sample were analysed in order to determine population means using the default software of the instrument.

### Mathematical modelling

The full model is described in detail in the Supplementary Material. The system of ordinary differential equations was implemented in MATLAB2019b (The MathWorks Inc., Natick, MA, USA) and its behaviour simulated using the in-built stiff solver *ode15s*. All simulations where initiated with 10 molecules of each protein species, including ribosomes, and 10^3^ molecules of energy. All simulations were run to steady state by increasing the simulation time span until the maximal absolute value of the derivate was very small (< 10^−3^). Isocost lines were simulated by incorporating two additional genes as described in the Supplementary Material.

### Model fitting

The model was parameterized using MATLAB’s in-built genetic algorithm function *ga*, with a large population of 320, and fitting ceasing when 100 generations pass without changing the cost function output significantly. The model was fit in the exponential growth regime and growth rate was varied by simulating the response of the model to a range of *ϕ*_*e*_ values. The cost function was made of up three sums of squared errors relating to the growth rate *σ*_*λ*._ and mass fraction of RNA polymerase *σ*_*P*_ and ribosomes *σ*_*R*_. *σ*_*λ*_ was defined as the sum of the squared error between the simulated growth rate *λ*_*sim*_ and the experimental growth rate *λ*_*exp*_:

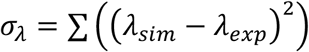

Given that the mass fraction of the RNA polymerase Φ_*p*_ and ribosomes Φ_*R*_ are on different scales of magnitude, the sum of squared error was normalized by the sum of the respective experimental data squared:

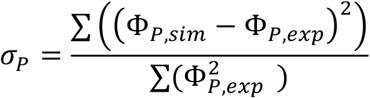

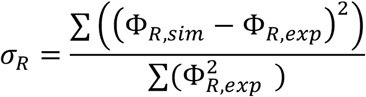

The cost function used for data fitting was:

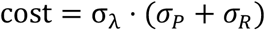

The parameters fit and their bounds are shown in the Supplementary Materials.

## Supporting information

Supplementary model description

## Acknowledgements

The authors would like to thank Dr Manuel Salvador for the support developing the chemostat and the insightful discussions on the liquid handling system. The authors are indebted to Dr Helen King, Dr Mandy Fivian-Hughes, Fiona Watt and Anita Sicilia for their technical assistance. JK and JIJ acknowledge the support received from the Biology and Biotechnology Research Council (grants BB/M009769/1 and BB/T011289/1 from the ERA-Cobiotech programme of the EU). APSD and DGB acknowledge the support of The Leverhulme Trust (grant RPG-2017-284).

