## Supplementary figures and methods for "The interplay between growth rate and nutrient quality defines gene expression capacity"

by

Juhyun Kim, Alexander P.S. Darlington, Declan G. Bates
and Jose I. Jimenez

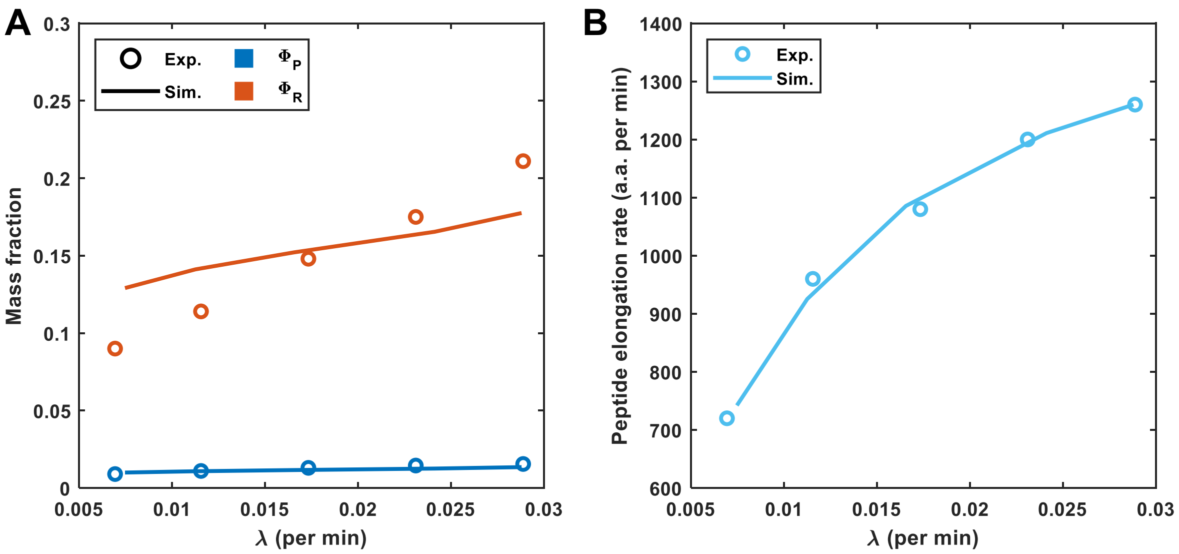


**Figure S1. Quality of data fitting. (A)** The model was fit as described in the main text. The simulated mass fraction of the RNA polymerase and ribosomes are shown with the experimental data from Bremer and Dennis (1996). **(B)** The peptide elongation values reported in the same publication were not used in the data fitting but simulations show good agreement.

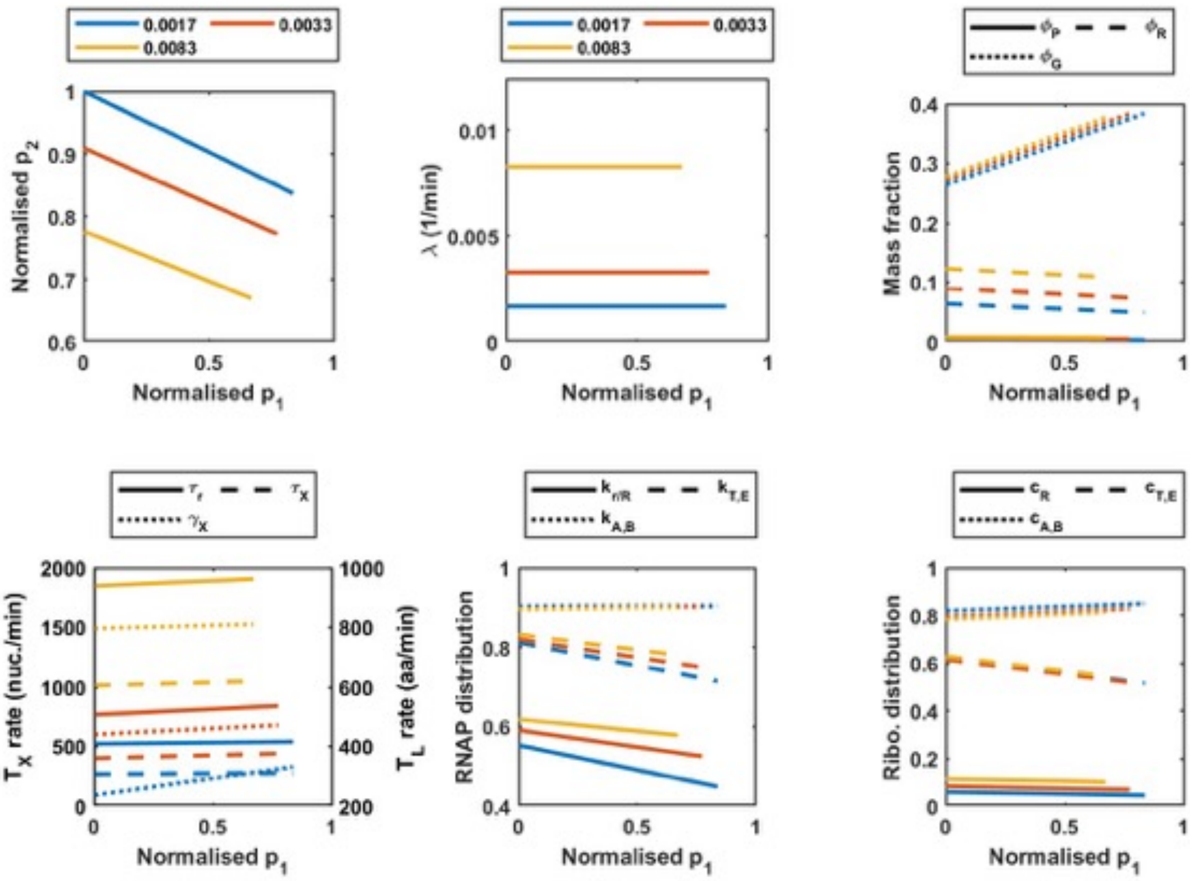


**Figure S2. Full simulation results from Figure 2.** Simulations of the steady state concentration of RFP and GFP, normalised by maximum protein production for a different growth rates. Simulations were carried out as described in the main text. **(A)** Protein output. **(B)** Growth rate. **(C)** The mass fraction of the RNA polymerase ($\Phi_P$), ribosomes ($\Phi_R$) and circuit proteins ($\Phi_G$). **(D)** The mRNA ($\tau_X$) and rRNA ($\tau_r$) transcription elongation rate (plotted on the left axis, $T_X$ rate) and peptide elongation rate ($\gamma_X$) (plotted on the right axis, $T_L$ rate). **(E)** Proportion of transcribing RNA polymerases transcribing rRNA and r-protein genes ($k_{r,R}$), enzymes ($k_{T,E}$) and circuit genes ($k_{A,B}$) **(F)** Proportion of translating ribosomes translating r-proteins ($c_R$), enzymes ($c_{T,E}$) and circuit genes ($c_{A,B}$).

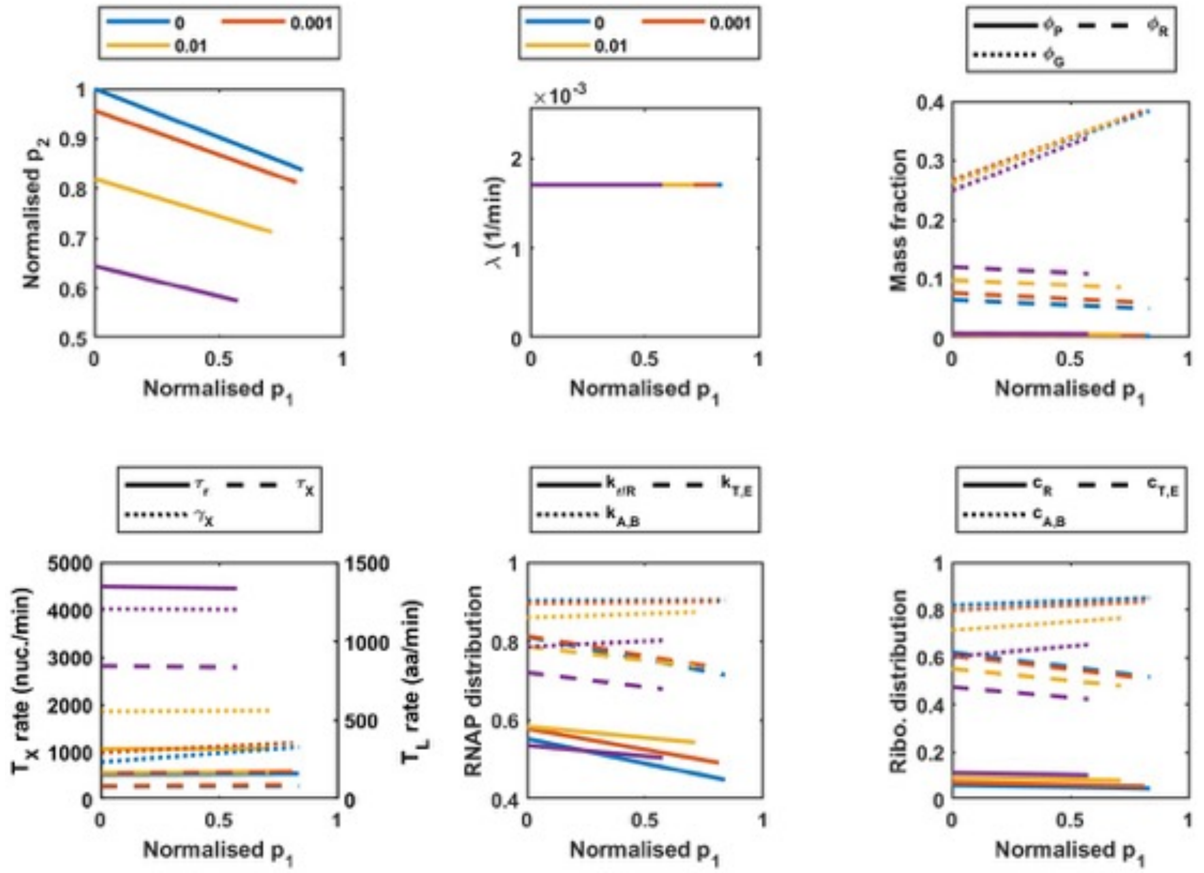


**Figure S3. Full simulation results from Figure 3.** Simulations of the steady state concentration of RFP and GFP, normalised by maximum protein production for a different RNA polymerase inhibition rates, $k_{rf}$. Simulations were carried out as described in the main text. **(A)** Protein output. **(B)** Growth rate. **(C)** The mass fraction of the RNA polymerase ($\Phi_P$), ribosomes ($\Phi_R$) and circuit proteins ($\Phi_G$). **(D)** The mRNA ($\tau_X$) and rRNA ($\tau_r$) transcription elongation rate (plotted on the left axis, $T_X$ rate) and peptide elongation rate ($\gamma_X$) (plotted on the right axis, $T_L$ rate). **(E)** Proportion of transcribing RNA polymerases transcribing rRNA and r-protein genes ($k_{r,R}$), enzymes ($k_{T,E}$) and circuit genes ($k_{A,B}$) **(F)** Proportion of translating ribosomes translating r-proteins ($c_R$), enzymes ($c_{T,E}$) and circuit genes ($c_{A,B}$).

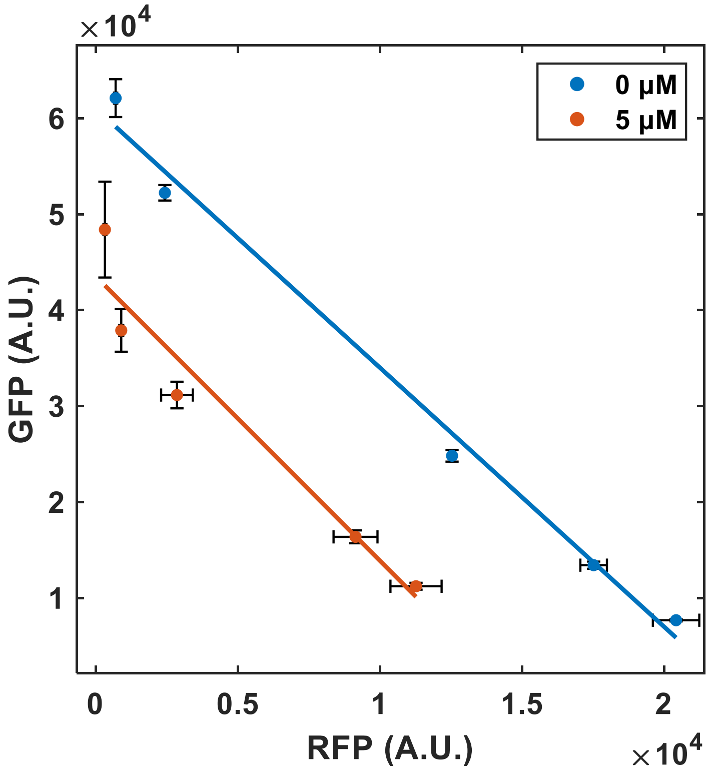


**Figure S4. Effect of selective inhibition of translation on the two-reporter circuit gene expression.** To characterize heterologous gene expression profile and the gene expression coupling under partial inhibition of translation, the dual reporter strain was grown in the chemostat at a set growth rate in the presence of a sublethal concentration of chloramphenicol (shown in the box) and different concentrations of AHL.

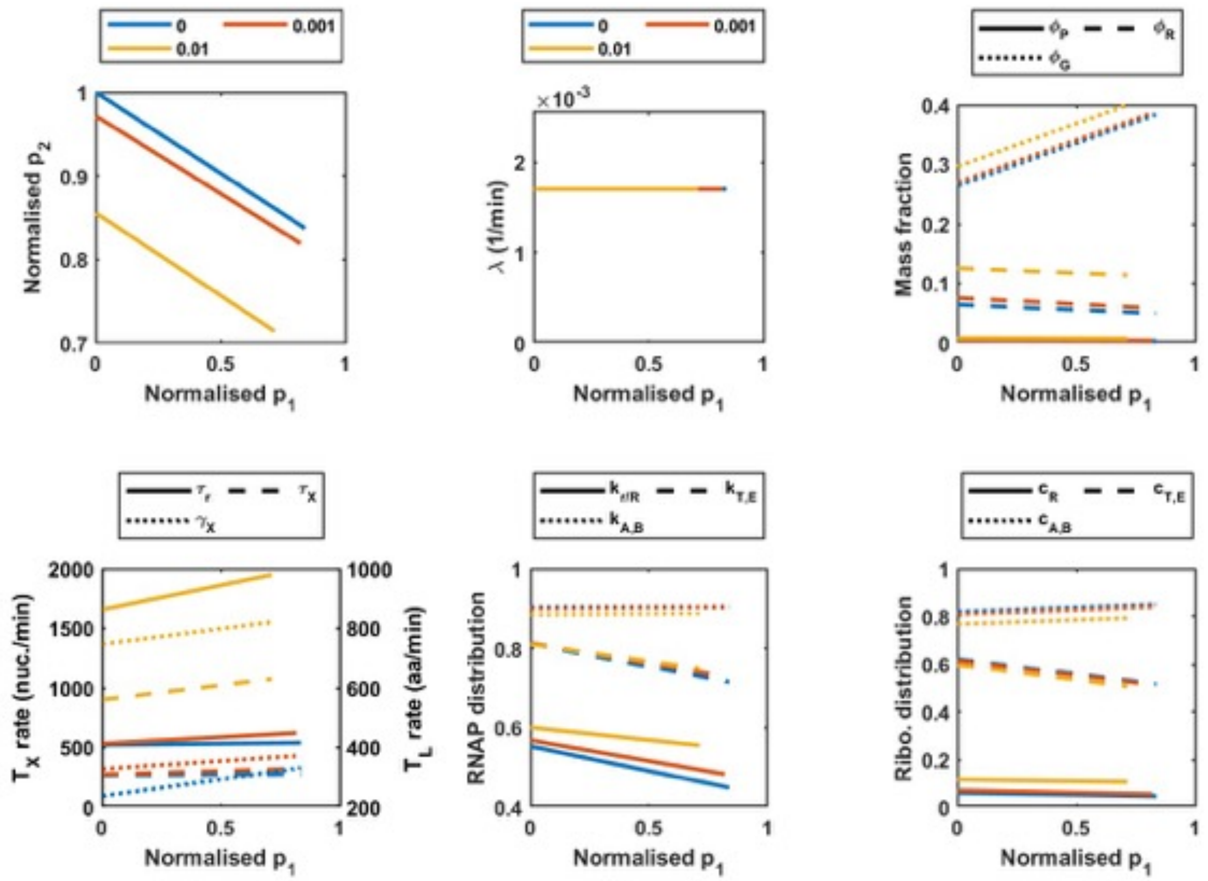


**Figure S5 Full simulation results for ribosome inhibition.** Simulations of the steady state concentration of RFP and GFP, normalised by maximum protein production for a different ribosome inhibition rates, $k_{cm}$. Simulations were carried out as described in the main text. **(A)** Protein output. **(B)** Growth rate. **(C)** The mass fraction of the RNA polymerase ($\Phi_P$), ribosomes ($\Phi_R$) and circuit proteins ($\Phi_G$). **(D)** The mRNA ($\tau_X$) and rRNA ($\tau_r$) transcription elongation rate (plotted on the left axis, $T_X$ rate) and peptide elongation rate ($\gamma_X$) (plotted on the right axis, $T_L$ rate). **(E)** Proportion of transcribing RNA polymerases transcribing rRNA and r-protein genes ($k_{r,R}$), enzymes ($k_{T,E}$) and circuit genes ($k_{A,B}$) **(F)** Proportion of translating ribosomes translating r-proteins ($c_R$), enzymes ($c_{T,E}$) and circuit genes ($c_{A,B}$).

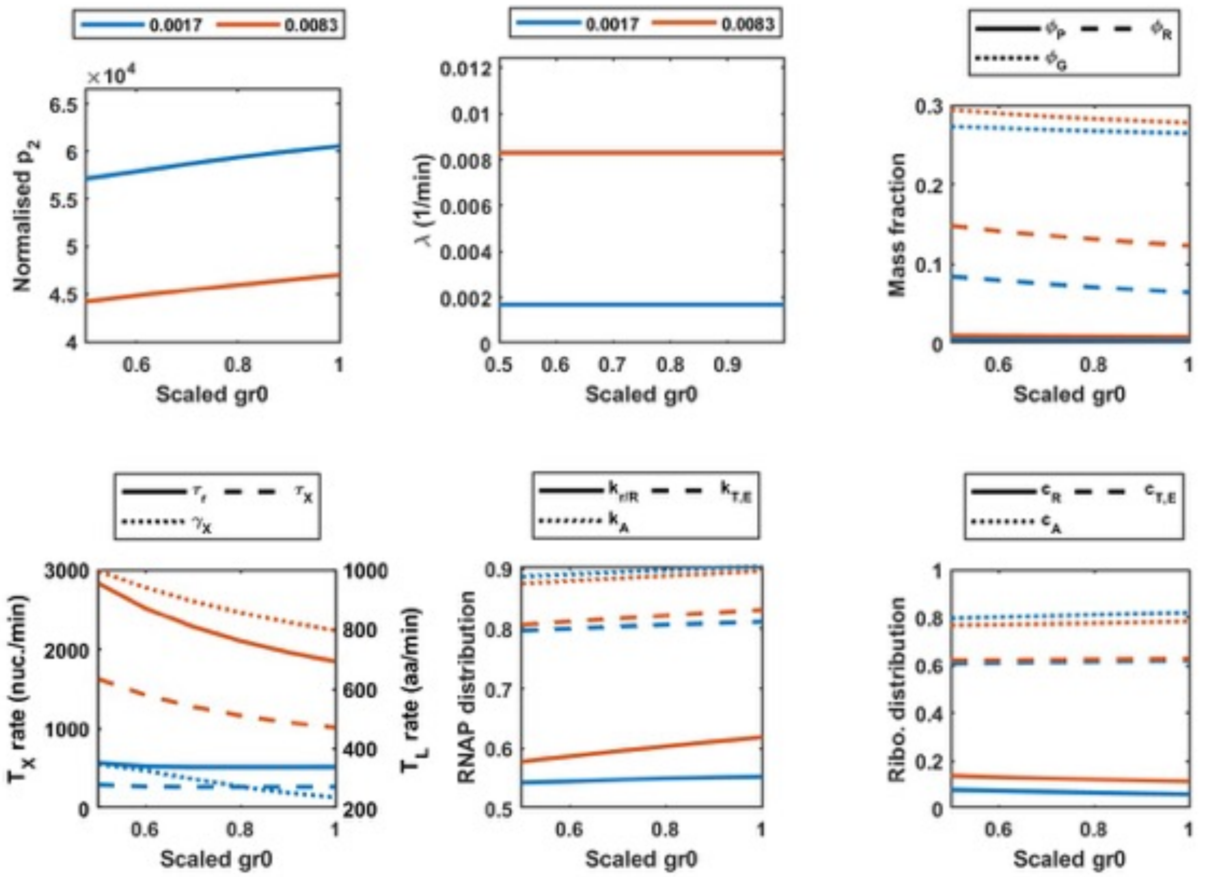


**Figure S6. Full simulation results from Figure 4.** Simulations of the steady state concentration of RFP and GFP for a different rRNA expression rates. Simulations were carried out as described in the main text. **(A)** Protein output. **(B)** Growth rate. **(C)** The mass fraction of the RNA polymerase ($\Phi_P$), ribosomes ($\Phi_R$) and circuit proteins ($\Phi_G$). **(D)** The mRNA ($\tau_X$) and rRNA ($\tau_r$) transcription elongation rate (plotted on the left axis, $T_X$ rate) and peptide elongation rate ($\gamma_X$) (plotted on the right axis, $T_L$ rate). **(E)** Proportion of transcribing RNA polymerases transcribing rRNA and r-protein genes ($k_{r,R}$), enzymes ($k_{T,E}$) and circuit genes ($k_{A,B}$) **(F)** Proportion of translating ribosomes translating r-proteins ($c_R$), enzymes ($c_{T,E}$) and circuit genes ($c_{A,B}$).

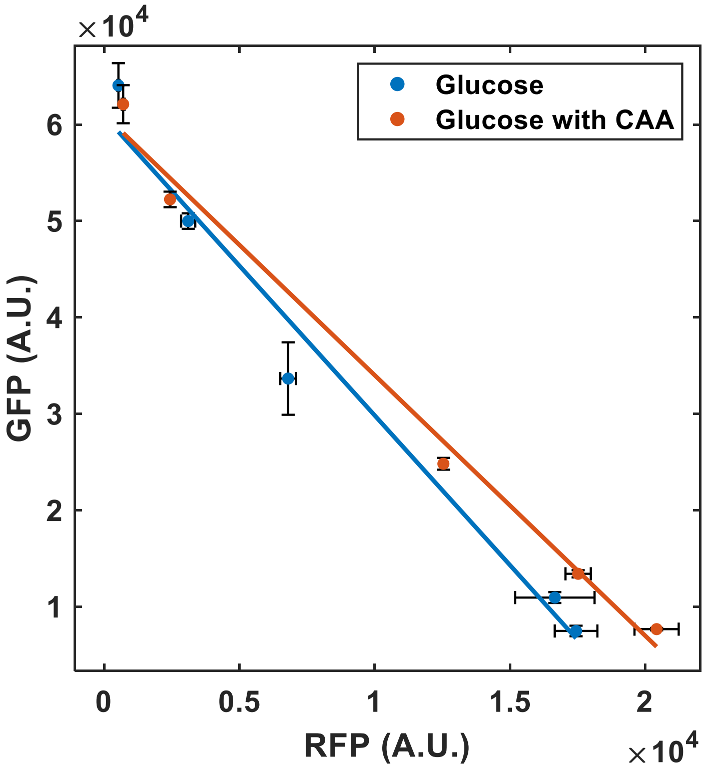


**Figure S7. Effect of additional nutrients on the circuit gene expressions.** The dual reporter strain was cultured in the continuous system containing M9 minimal medium supplemented with glucose as a sole carbon source. To determine the level of the gene expression coupling, different concentration of AHL applied into each bioreactor and measured both GFP and RFP intensities. The counterpart expression profile from Figure 2A also displayed in this figure.

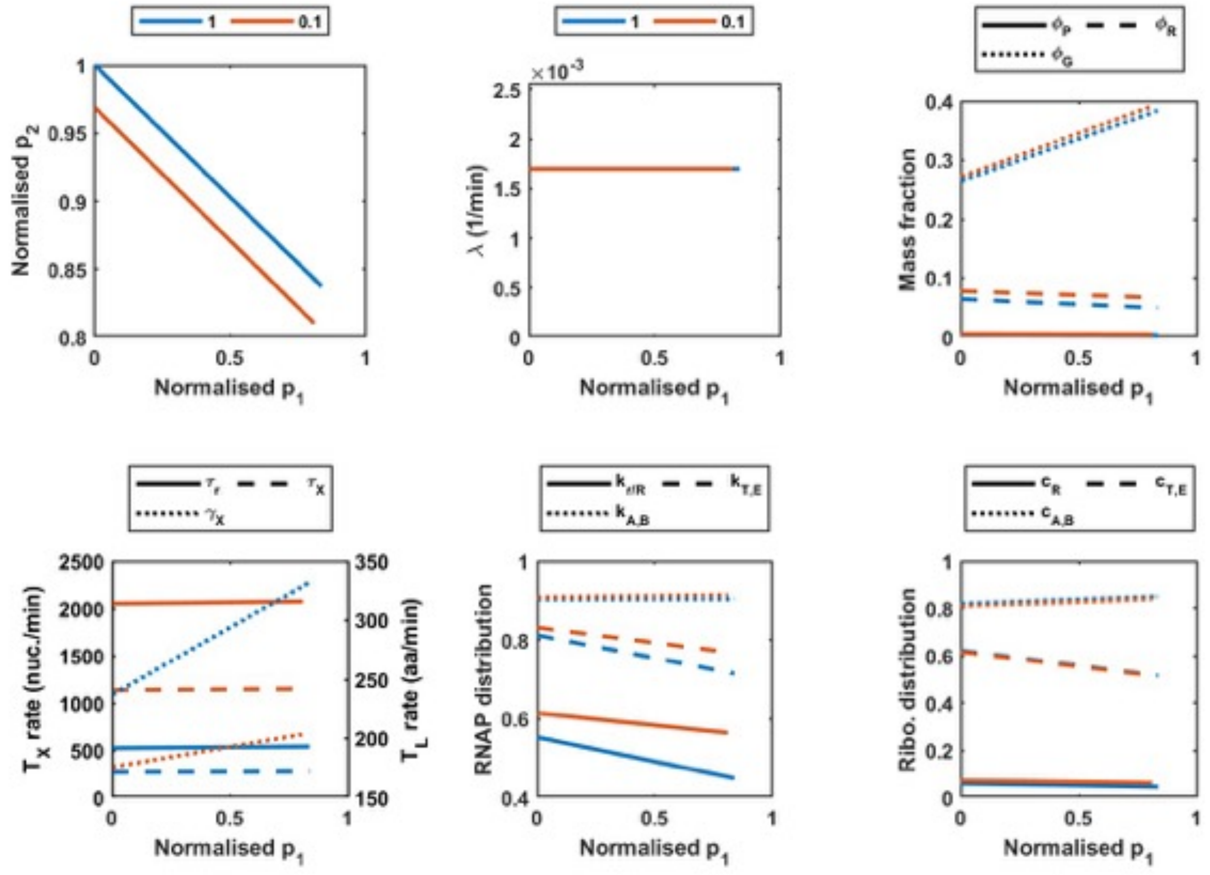


**Figure S8. Full simulation results from Figure 5.** Simulations of the steady state concentration of RFP and GFP for a different $\phi_a$ values. Simulations were carried out as described in the main text. **(A)** Protein output. **(B)** Growth rate. **(C)** The mass fraction of the RNA polymerase ($\Phi_P$), ribosomes ($\Phi_R$) and circuit proteins ($\Phi_G$). **(D)** The mRNA ($\tau_X$) and rRNA ($\tau_r$) transcription elongation rate (plotted on the left axis, $T_X$ rate) and peptide elongation rate ($\gamma_X$) (plotted on the right axis, $T_L$ rate). **(E)** Proportion of transcribing RNA polymerases transcribing rRNA and r-protein genes ($k_{r,R}$), enzymes ($k_{T,E}$) and circuit genes ($k_{A,B}$) **(F)** Proportion of translating ribosomes translating r-proteins ($c_R$), enzymes ($c_{T,E}$) and circuit genes ($c_{A,B}$).

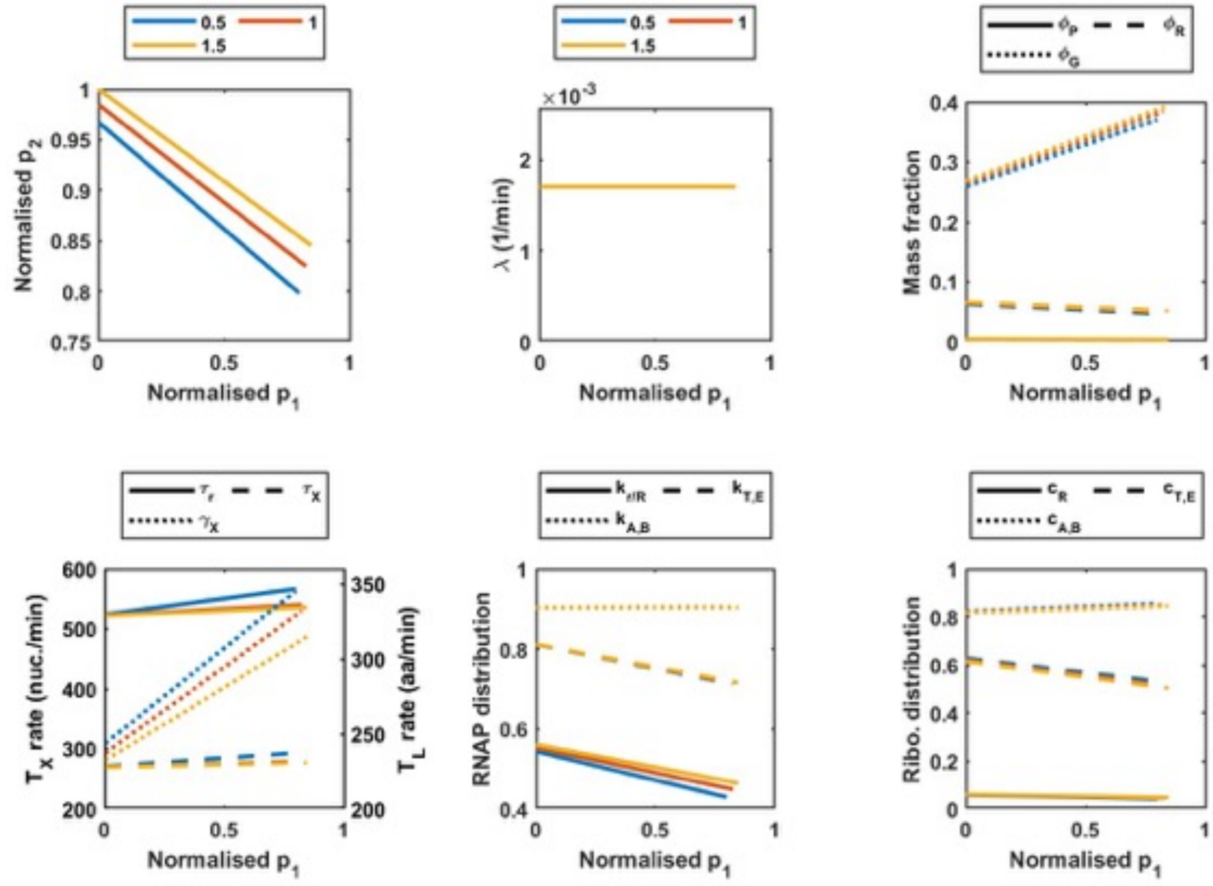


**Figure S9. Full simulation results for varying $o_X$.** Simulations of the steady state concentration of RFP and GFP for a 0.5, 1 and 1.5 times the nominal value of $o_X$. Simulations were carried out as described in the main text. **(A)** Protein output. **(B)** Growth rate. **(C)** The mass fraction of the RNA polymerase ($\Phi_P$), ribosomes ($\Phi_R$) and circuit proteins ($\Phi_G$). **(D)** The mRNA ($\tau_X$) and rRNA ($\tau_r$) transcription elongation rate (plotted on the left axis, $T_X$ rate) and peptide elongation rate ($\gamma_X$) (plotted on the right axis, $T_L$ rate). **(E)** Proportion of transcribing RNA polymerases transcribing rRNA and r-protein genes ($k_{r,R}$), enzymes ($k_{T,E}$) and circuit genes ($k_{A,B}$) **(F)** Proportion of translating ribosomes translating r-proteins ($c_R$), enzymes ($c_{T,E}$) and circuit genes ($c_{A,B}$).

**Table S1. Bacterial strains, plasmids and oligonucleotides used in this study**

| ***E. coli* strains** | **Description** | **Reference** |
|---|---|---|
| MG1655 | $F^-$, $\lambda^-$, *ilvG*$^-$, *rfb-50*, *rph-1* | (Blattner *et al*, 1997) |
| MG1655 L9-*msf*GFP | MG1655 derivative with a C-terminal msf GFP tag in 50S ribosomal protein L9, encoded by *rplI* gene | This study |
| BW25113 | Parental strain for the Keio Collection of single gene knockouts Δ(*araD-araB*)567,Δ*lacZ4787*(::*rrnB-3*), λ-, *rph-1*, Δ(*rhaD-rhaB*)568, *hsdR514* | (Baba *et al*, 2006) |
| BW25113 (Δ relA) | BW25113 derivate with a deletion of *relA* genes | Keio collection |
| BW25113 (Δ spoT) | BW25113 derivate with a deletion of *spoT* genes | This study |
| DH5α | Cloning host: $F^-$ Φ80 *lacZ*Δ*M*Δ15 (*lacZYA-argF*), U169, *recA1, endA1, hsdR17, R-M+, supE44, thil, gyrA, relA1* | (Hanahan & Meselson, 1983) |
| DH5α λpir | DH5α λpir phage lysogen | Victor de Lorenzo's collection |
| SQ37 | MG1655 derivate, Δ*rrnE* | (Quan *et al*, 2015) |
| SQ40 | MG1655 derivate, Δ*rrnEG* | (Quan *et al.*, 2015) |
| SQ49 | MG1655 derivate, Δ*rrnGBA* | (Quan *et al.*, 2015) |
| SQ53 | MG1655 derivate, Δ*rrnGBAD* | (Quan *et al.*, 2015) |
| SQ78 | MG1655 derivate, Δ*rrnGADE* | (Quan *et al.*, 2015) |
| **Plasmid** | **Description** | **Reference** |
| pEMG | Suicide plasmid, $Km^R$, oriR6K, *lacZa* with two flanking I-SceI sites | (Martinez-Garcia & de Lorenzo, 2011) |
| pEMG-rplI-msfGFP | Same as pEMG but carrying the msf GFP gene with upstream and downstream flanking regions of the C-terminus of *E. coli* rplI gene | This study |
| pKD4 | Template for Km cassette, oriR6Kgamma, *bla, aphA* | (Datsenko & Wanner, 2000) |
| pKD46 | Red recombinase expression vector, repA101ts, oriR101, $P_{araB}$ *exo, bet, gam araC bla* | (Datsenko & Wanner, 2000) |
| pSEVA63-Dual | pSEVA631 carrying the circuit MBP 1.0 | (Darlington *et al*, 2018) |

| Oligos | Sequence (5' --> 3') |
|---|---|
| TS1$^{rplI}$F | AGGGATAACAGGGTAATCTGCGCTCGCTACCTGTCCCTGCT |
| TS1$^{rplI}$R | TTTACTGCCACCGCCACCGCTTTCAGCTACTACGTTTACGA |
| rplI-msfGFP-F | GAAAGCGGTGGCGGTGGCAGTAAAGGTGAAGAACTGTTCACCG |
| rplI-msfGFP-R | TACGTCTCGTTGAATAACGAATTATTTGTAGAGTTCATCCAT |
| TS2$^{rplI}$F | CATGGATGAACTCTACAAATAATTCGTTATTCAACGAGACGT |
| TS2$^{rplI}$R | GCCTGCAGGTCGACTCTAGAGTATTTATTGCAAGATGTCGAAT |
| spoT KO F | TTACCGCTATTGCTGAAGGTCGTCGTTAATCACAAAGCGGGTCGC CCTTGGTGTAGGCTGGAGCTGCTTC |
| spoT KO R | CGTGCATAACGTGTTGGGTTCATAAAACATTAATTTCGGTTTCGG GTGACATGGGAATTAGCCATGGTCC |
