## Supplementary model description for "The interplay between growth rate and nutrient quality defines gene expression capacity"

### 1 Model description

This model builds on the trade-off models presented in [1, 2] and aims to incorporate the additional competition for RNA polymerase and a more complex metabolism. The model couples mechanistic models of transcription and translation with phenomenological descriptions of microbial growth and a simple representation of metabolism to produce a simple tractable model capable of recapitulating the different distributions of cellular proteins (such as transporters, enzymes, host factors, RNA polymerase, ribosomes and circuit genes) in addition to partitions of RNAP for transcription of r or mRNA coding genes. Applying the Law of Mass Action to the reactions listed below a single host model of 27 coupled ordinary differential equations capturing the following universal constraints:

(i) Finite biosynthetic capacity provided by a simple metabolism

- transport and metabolic enzymes create limitation in nutrient update
- external nutrients can also be constrained allowing modelling of (i) mid-exponential growth, (ii) batch culture and (iii) chemostat.

(ii) Competition for total number of proteins (‘space’)

(iii) Competition for RNA polymerases by mRNA and rRNA promoters

- including increasing copy number with growth rate

(iv) Competition for ribosomes by mRNAs

(v) Resource biosynthesis

- including autocatalysis – i.e. gene expression resources are required for their own production
- including the underlying metabolic limitations on reaction rates

(vi) Dynamic feedback between growth rate and resource biosynthesis

### 2 Model ODEs

#### 2.1 Metabolism

We consider growth in the presence of an external substrate $s_x$ which is imported into the cell by transporter proteins $p_T$ to produce $s_i$ which itself is converted into a proxy ‘energy’ species $e$ and amino acids $a$ or nucleotides $n$ by enzymes $p_E$. Considering metabolism and growth in this way allows modelling of:

(i) Single cells during exponential growth (where the external resource is constant and population dynamics are ignored, i.e. $dN/dt$ and $ds_x/dt$ is set to zero)

(ii) Batch culture (where the external resources deplete and the population grows over time, where $k_{in} = \delta = 0$)

(iii) Growth in a chemostat (where at steady state $dN/dt = 0$ when the growth rate $\lambda$ equals the dilution rate $\delta$)

This allows us to replicate the multiple culture methods available which the preliminary data suggests that coupling is dependent upon culture condition. The dynamics of the external and internal substrates are:

$$\frac{ds_x}{dt} = \begin{cases} \underbrace{0}_{\text{constant ext. substrate}} & \text{Exponential growth} \\ \underbrace{-\left(\frac{v_T \cdot s_x \cdot p_T}{\kappa_T + s_x}\right) \cdot N}_{\text{consumption of } s_x \text{ by } N \text{ bacteria}} & \text{Batch growth} \\ \underbrace{k_{in}}_{s_x \text{ influx}} - \underbrace{\left(\frac{v_T \cdot s_x \cdot p_T}{\kappa_T + s_x}\right) \cdot N}_{\text{consumption of } s_x \text{ by } N \text{ bacteria}} - \underbrace{\delta \cdot s_x}_{s_x \text{ efflux}} & \text{Chemostat growth} \end{cases} \tag{1}$$

$$\frac{ds_i}{dt} = \underbrace{\left(\frac{v_T \cdot s_x \cdot p_T}{\kappa_T + s_x}\right)}_{\text{substrate import}} - \underbrace{\left(\frac{v_E \cdot s_i \cdot p_E}{\kappa_E + s_i}\right)}_{\text{converston of } s_i \text{ to } e} - \underbrace{2 \cdot \left(\frac{v_E \cdot s_i \cdot e \cdot p_E}{{\kappa_E}^2 + \kappa_E \cdot s_i + \kappa_E \cdot e + s_i \cdot e}\right)}_{\text{converston of } s_i \text{ to } a \text{ and } n} - \underbrace{\lambda \cdot s_i}_{\text{dilution}} \tag{2}$$

The internal 'energy' intermediate $e$ is produced from the internalised substrate by the metabolic enzymes $p_E$. The incorporation of each nucleotide into an rRNA or mRNA utilises $\phi_X$ energy molecules - this approximates the energy required for the production of nucleotides and for joining nucleotides during elongation etc. – we set $\phi_X$ to 10. The incorporation of each amino acid into the nascent peptide change utilises $\phi_L$ energy molecules - this approximates tRNA charging and peptide bond formation etc. – we set $\phi_L$ to 6. We approximate the carbon and energy requirements of nucleotide or amino acid biosynthesis by considering the production of these pools from both $s_i$ and $e$ catalysed by the enzyme fraction $p_E$.

The dynamics of the internal substrates are:

$$\begin{aligned} \frac{de}{dt} = & \underbrace{\phi_e \cdot \left(\frac{v_E \cdot s_i \cdot p_E}{\kappa_E + s_i}\right)}_{\text{substrate conversion to } e} - \underbrace{\lambda \cdot e}_{dilution} \ldots \\ & \ldots - \underbrace{\phi_X \cdot \left(n_r \cdot T_X(k_r,\ e) + \sum_i \left(3 \cdot n_i \cdot T_X(k_i,\ e)\right)\right)}_{\text{energy consumption by transcription}} - \underbrace{\phi_L \cdot \sum_i \left(n_i \cdot T_L(c_i,\ e,\ a)\right)}_{\text{energy consumption by translation}} \ldots \\ & \ldots - \underbrace{2 \cdot \left(\frac{v_E \cdot s_i \cdot e \cdot p_E}{{\kappa_E}^2 + \kappa_E \cdot s_i + \kappa_E \cdot e + s_i \cdot e}\right)}_{\text{converston of } s_i \text{ and } e \text{ to } a \text{ and } n} \end{aligned} \tag{3}$$

$$\frac{dn}{dt} = \underbrace{\phi_n \cdot \left(\frac{v_E \cdot s_i \cdot e \cdot p_E}{{\kappa_E}^2 + \kappa_E \cdot s_i + \kappa_E \cdot e + s_i \cdot e}\right)}_{\text{production of } n \text{ from } s_i \text{ and } e} - \underbrace{\left(n_r \cdot T_X(k_r,\ e) + \sum_i \left(3 \cdot n_i \cdot T_X(k_i,\ e)\right)\right)}_{\text{consumption by transcription}} - \underbrace{\lambda \cdot n}_{dilution} \tag{4}$$

$$\frac{da}{dt} = \underbrace{\phi_a \cdot \left(\frac{v_E \cdot s_i \cdot e \cdot p_E}{{\kappa_E}^2 + \kappa_E \cdot s_i + \kappa_E \cdot e + s_i \cdot e}\right)}_{\text{production of } a \text{ from } s_i \text{ and } e} - \underbrace{\sum_i \left(n_i \cdot T_L(c_i,\ e,\ a)\right)}_{\text{consumption by translation}} - \underbrace{\lambda \cdot a}_{dilution} \tag{5}$$

#### 2.2 Gene expression model

**Accounting for gene copy number changes.** DNA replication is faster than cell division and so, especially at higher growth rates, the genome exists in multiple copies per cell. We describe the copy number phenomenologically by relating it to growth rate:

$$g_{X,T} = g_{X,0} \cdot \exp(q_X \cdot \lambda) \tag{6}$$

where $g_{X,0}$ is the copy number present in the chromosome is at one copy per cell and $q_X$ controlling the steepness of the association.

A gene's promoter can be either free ($g$), occupied by an RNAP polymerase ($k$) or, in the presents of rifampacin, the later can be in a 'stalled/sequestered' state ($y$, see Section 2.6). Therefore at each point in time:

$$g_{X,T} = g_X + k_X + y_X \qquad \therefore \qquad g_X = g_{X,0} \cdot \exp(q_X \cdot \lambda) - k_X - y_X \tag{7}$$

**Gene expression model.** The free promoter for a host protein-encoding gene $g_j$ is bound by the free RNA polymerase $P$ at maximal rate $\beta_j$ to form the transcription complex $k_j$. The maximal rate of binding $\beta_j$ is scaled by the regulatory function $\mathcal{R}_j(\cdot)$, see below. The RNA polymerase unbinds at rate $\mu_j$. The transcription complex is 'consumed' by transcription $T_X(\cdot)$. The messenger RNA $m_j$ is produced by transcription $T_X$ and is bound by ribosomes $R$ at rate $b_j$ to produce the translational complex $c_j$. The ribosome unbinds at rate $u_j$. The mRNA is released upon termination of translation at rate $T_L(\cdot)$. Translation produces ]proteins $p_j$. All species dilute due to the growth while mRNA have an additional decay rate $\delta_{m_j}$. The dynamics of gene expression are:

$$\frac{dk_j}{dt} = -\underbrace{\lambda \cdot k_j}_{\text{dilution}} + \underbrace{\mathcal{R}_j(\cdot) \cdot \beta_j \cdot g_j \cdot P}_{\text{RNAP binding}} - \underbrace{\mu_j \cdot k_j}_{\text{RNAP unbinding}} - \underbrace{T_X(k_j,\ e)}_{\text{transcription}} \qquad {\scriptstyle j \in \{T,\ E,\ H,\ P,\ R,\ r\}} \tag{8}$$

$$\frac{dm_j}{dt} = -\underbrace{(\lambda + \delta_{m_j}) \cdot m_j}_{\text{dilution/decay}} + \underbrace{T_X(k_j,\ e)}_{\text{transcription}} + \underbrace{T_L(c_j,\ e,\ a)}_{\text{translation}} - \underbrace{b_j \cdot m_j \cdot R}_{\text{ribo. binding}} + \underbrace{u_j \cdot c_j}_{\text{ribo. unbinging}} \qquad {\scriptstyle j \in \{T,\ E,\ H,\ P,\ R\}} \tag{9}$$

$$\frac{dc_j}{dt} = -\underbrace{\lambda \cdot c_j}_{\text{dilution}} + \underbrace{b_j \cdot m_j \cdot R}_{\text{ribo. binding}} - \underbrace{u_j \cdot c_j}_{\text{ribo. unbinging}} - \underbrace{T_L(c_j,\ e,\ a)}_{\text{translation}} \qquad {\scriptstyle j \in \{T,\ E,\ H,\ P,\ R\}} \tag{10}$$

$$\frac{dp_j}{dt} = -\underbrace{\lambda \cdot p_j}_{\text{dilution}} + \underbrace{T_L(c_j,\ e,\ a)}_{\text{translation}} \qquad {\scriptstyle j \in \{T,\ E,\ H\}} \tag{11}$$

**Regulating gene expression.** The maximal rate of binding of the the RNA polymerase to the promoter is scaled by the regulatory function $\mathcal{R}_j(\cdot)$ which scales by the cell's internal energy status ($e/(o+e)$). The amino acid and host protein fraction are also subject to feedback inhibition:

$$\mathcal{R}_j = \frac{e}{o_j + e} \tag{12}$$

$$\mathcal{R}_H = \left(\frac{e}{o_H + e}\right) \cdot \left(\frac{1}{1 + (p_H/\kappa_H)^{h_H}}\right) \tag{13}$$

We assume that all gene expression resources respond to the cell's energy state in the same way (i.e. $o_r = o_R = o_P$) and we assume that all other genes (including circuit genes) respond to the cell's energy state in the same way (i.e. $o_T = o_E = o_H$), as suggested by the observations in [3].

##### 2.3 Resource biosynthesis

**RNA polymerase dynamics.** $dk_P/dt$, $dm_P/dt$ and $dc_X/dt$ follow the dynamics outlined in Eq. 8 to 10. The dynamics of the RNA polymerase protein itself are given by:

$$\frac{dP}{dt} = \underbrace{-\underbrace{\lambda \cdot P}_{\text{dilution}} + \underbrace{T_L(c_P,\ e,\ a)}_{\text{translation}}}_{\text{protein production and decay}} + \underbrace{\sum_j \Big(T_X(k_j,\ e) - \mathcal{R}_j(\cdot) \cdot \beta_j \cdot P \cdot g_j + \mu_j \cdot k_j\Big)}_{\text{gene } g_j \text{ expression dynamics, } j \in \{T,\ E,\ A,\ H,\ P,\ R,\ r\}} \tag{14}$$

**rRNA ODEs** The RNAP-rRNA gene dynamics ($g_r$ and $dk_r/dt$) follow the same dynamics outlined in Eq. 7 and 8. The transcribed rRNA $r$ binds to 'empty' ribosomes $p_R$ at rate $b_\rho$ to produce functional ribosomes $R$. The functional ribosome can dissociate at rate $u_\rho$. Note the rRNA dilutes but does not decay unlike mRNAs. The protein component of the ribosome (r-proteins) are expressed in the same way as other genes, therefore $dm_R/dt$, $dk_R/dt$, $dm_R/dt$ and $dc_R/dt$ follow the dynamics of Eq. 8 to 10. Both rRNA and r-proteins dilute due to growth (note that the rRNA does not degrade like mRNAs do). The dynamics of the rRNA and r-proteins are:

$$\frac{dr}{dt} = \underbrace{-\underbrace{\lambda \cdot r}_{\text{dilution}} + \underbrace{T_X(k_r,\ e)}_{\text{transcription}}}_{\text{rRNA production/dilution}} \underbrace{-\underbrace{b_\rho \cdot p_R \cdot r}_{\text{ribo. formation}} + \underbrace{u_\rho \cdot R}_{\text{ribo. dissociation}}}_{\text{functional ribosome production}} \tag{15}$$

$$\frac{dp_R}{dt} = \underbrace{-\underbrace{\lambda \cdot p_R}_{\text{dilution}} + \underbrace{T_L(c_R,\ e,\ a)}_{\text{translation}}}_{\text{r-protein production/dilution}} \underbrace{-\underbrace{b_\rho \cdot p_R \cdot r}_{\text{ribo. formation}} + \underbrace{u_\rho \cdot R}_{\text{ribo. dissociation}}}_{\text{functional ribosome production}} \tag{16}$$

**Ribosome dynamics.** The dynamics of the free functional ribosome follows:

$$\frac{dR}{dt} = \underbrace{\underbrace{-\lambda \cdot R}_{\text{dilution}} + \underbrace{+\, b_\rho \cdot p_R \cdot r - u_\rho \cdot R}_{\text{r-protein/rRNA binding dynamics}}}_{\text{free ribosome production and decay}} + \underbrace{\sum_j \Big(T_L(c_j,\ e,\ a) - b_j \cdot R \cdot m_j + u_j \cdot c_j\Big)}_{\text{mRNA } m_j \text{ expression dynamics, } j \in \{T,\ E,\ A,\ H,\ P,\ R\}} \tag{17}$$

##### 2.4 Population

The dynamics of the population depend on the culture conditions are given by:

$$\frac{dN}{dt} = \begin{cases} 0 & \text{Exponential growth} \\ \lambda \cdot N & \text{Batch growth} \\ (\lambda - \delta) \cdot N & \text{Chemostat growth} \end{cases} \tag{18}$$

##### 2.5 Transcription, translation and growth rate functions

In [1], the global translation rate is derived according to the principles in [4] and this conforms to a Michaelis-Menten like expression with a maximal rate scaled by substrate concentration with the same of curve determined by a 'threshold constant'.

Here, we assume the global translation rate is dependent upon both energy and amino acid concentration.

$$\gamma(e,\ a) = \gamma_{max} \cdot \left(\frac{e}{\kappa_{\gamma,\ e} + e}\right) \cdot \left(\frac{a}{\kappa_{\gamma,\ a} + a}\right) \tag{19}$$

We assume that the energy threshold $\kappa_{\gamma,\ e}$ is equal to the amino threshold $\kappa_{\gamma,\ a}$; i.e. $\kappa_\gamma = \kappa_{\gamma,\ e} = \kappa_{\gamma,\ a}$. Therefore the global translation rate is given by:

$$\gamma(e,\ a) = \frac{\gamma_{max} \cdot e \cdot a}{{\kappa_\gamma}^2 + \kappa_\gamma \cdot e + \kappa_\gamma \cdot a + e \cdot a} \tag{20}$$

Protein production from translation complex $c_j$ is given by the global translation rate $\gamma$ scaled by the length of the protein in amino acids $n_j$:

$$T_L(c_j,\ e,\ a) = \frac{\gamma(e,\ a)}{n_j} \cdot c_j \tag{21}$$

We apply the same assumptions to transcription therefore global rRNA, $\tau_r$, and mRNA, $\tau_X$, elongation rates are given by:

$$\tau_r(e,\ n) = \frac{\tau_{r,max} \cdot e \cdot n}{{\kappa_{\tau,r}}^2 + \kappa_{\tau,r} \cdot e + \kappa_{\tau,r} \cdot n + e \cdot n} \qquad \tau_X(e,\ n) = \frac{\tau_{X,max} \cdot e \cdot n}{{\kappa_{\tau,X}}^2 + \kappa_{\tau,X} \cdot e + \kappa_{\tau,X} \cdot n + e \cdot n} \tag{22}$$

The rate of mRNA production $T_X(\cdot)$ from the transcription complex, $k_j$, is the global rate $\tau_X$ scaled by length of the mRNA for gene $j$ (i.e. three times their length in amino acids $n_j$):

$$T_X(k_j,\ e,\ n) = \frac{\tau_X(e,\ n)}{3 \cdot n_j} \cdot k_j \tag{23}$$

The rate of rRNA production from the transcription complex $k_r$ is calculated similarly:

$$T_X(k_r,\ e,\ n) = \frac{\tau_r(e,\ n)}{n_r} \cdot k_r \tag{24}$$

(note that the length of the rRNA $n_r$ is measured in nucleotides not codons).

The growth rate is proportional to the global translation rate:

$$\lambda = (1/M_0) \cdot \gamma(e,\ a) \cdot \sum_j (c_j) \tag{25}$$

#### 2.6 Accounting for antibiotic inhibition

Both rifampacin and chloramphenicol inhibit elongation. Consistent with the method in [1], we model this inhibition as a sequestration of transcription or translation complexes ($k_j$ and $c_j$ respectively) to an inert state ($y_j$ and $z_j$ respectively).

We modify Eq. 8 to account for this sequestration and include an additional equation for the inhibited complex. *N.B.* that we assume this is a non-reversible reaction. For all genes $j \in \{T,\ E,\ H,\ P,\ R,\ r\}$

$$\frac{dk_j}{dt} = -\underbrace{\lambda \cdot k_j}_{\text{dilution}} + \underbrace{R_j \cdot \beta_j \cdot g_j \cdot P}_{\text{RNAP binding}} - \underbrace{\mu_j \cdot k_j}_{\text{RNAP unbinding}} - \underbrace{T_X(k_j,\ e)}_{\text{transcription}} - \underbrace{k_{rf} \cdot k_j}_{\text{rifampicin action}} \tag{26}$$

$$\frac{dy_j}{dt} = -\underbrace{\lambda \cdot y_j}_{\text{dilution}} + \underbrace{k_{rf} \cdot k_j}_{\text{rifampicin action}} \tag{27}$$

Similarity, Eq. 10 is modified and the inhibited complex added for protein encoding genes ($X \in \{T,\ E,\ H,\ P,\ R\}$):

$$\frac{dc_j}{dt} = -\underbrace{\lambda \cdot c_j}_{\text{dilution}} + \underbrace{b_j \cdot m_j \cdot R}_{\text{ribo. binding}} - \underbrace{u_j \cdot c_j}_{\text{ribo. unbinging}} - \underbrace{T_L(c_j,\ e,\ a)}_{\text{translation}} - \underbrace{k_{cm} \cdot c_j}_{\text{chloramphenicol action}} \tag{28}$$

$$\frac{dz_j}{dt} = -\underbrace{\lambda \cdot z_j}_{\text{dilution}} + \underbrace{k_{cm} \cdot c_j}_{\text{rifampicin action}} \tag{29}$$

### 3 Computational methods

#### 3.1 Simulating the model modelling

The system of ordinary differential equations describing the full model was implemented in MATLAB2019b (The MathWorks Inc., Natick, MA, USA) and its behaviour simulated using the in-built stiff solver *ode15s*. All simulations where initiated with 10 molecules of each protein species, including ribosomes, and 103 molecules of energy. All simulations were run to steady state by increasing the simulation time span until the maximal absolute value of the derivate was small ($< 10^{-3}$ molecules per min). Isocost lines were simulated by incorporating two additional genes as described below.

#### 3.2 Model fitting

The model was parameterized using MATLAB's in built genetic algorithm function (*ga*, with a large population of 320, with a fitting ceasing when 100 generations pass without changing in cost function output significantly). The model was fit in the exponential growth regime and growth rate was varied by simulating the response of the model to a range of $\phi_e$ values. The cost function was made of up three sums of squared errors relating to the growth rate $\sigma_\lambda$ and mass fraction of RNA polymerase $\sigma_P$ and ribosomes $\sigma_R$. $\sigma_\lambda$ was defined as the sum of the squared error between the simulated growth rate $\lambda_{sim}$ and the experimental growth rate $\lambda_{exp}$:

$$\sigma_\lambda = \sum \left( (\lambda_{sim} - \lambda_{exp})^2 \right) \tag{30}$$

Given that the mass fraction of the RNA polymerase $\Phi_P$ and ribosomes $\Phi_R$ are on different scales of magnitude the sum of squared error was normalised by the sum of the respective experimental data squared:

$$\sigma_P = \frac{\sum \left( (\Phi_{P,sim} - \Phi_{P,exp})^2 \right)}{\sum ({\Phi_{P,exp}}^2)} \qquad \sigma_R = \frac{\sum \left( (\Phi_{R,sim} - \Phi_{R,exp})^2 \right)}{\sum ({\Phi_{R,exp}}^2)} \tag{31}$$

The cost function used for data fitting was:

$$cost = \sigma_\lambda \cdot (\sigma_P + \sigma_R) \tag{32}$$

The parameters fit and their bounds are shown in Supplementary Table 1. Known parameters are shown in Supplementary Table 2.

#### 3.3 Simulating circuit gene expression

We introduce circuit genes by introducing new species and equations describing the protein of transcription complexes, mRNA, translation complexes and proteins. These equations show the same dynamics as shown in Section 2.2, and their antibiotic inhibited complexes show the same dynamics described in Section 2.6. We also modify Equations 3, 5 and 4, which describe the dynamics of the energy species, amino acids and nucleotides respectively, to take account of the additional demand for metabolites. We also modify the dynamics of the

Table 1: **Parameter values identified during data fitting**

| Parameter | Units | L.B. | U.B. | Fit value | Notes |
|---|---|---|---|---|---|
| min $\phi_e$ | unitless | $-3$ | 2 | $-0.26496$ | Varied on a log scale $10^x$, $x$ is fit. $\phi_e = 0.543$ |
| max $\phi_e$ | unitless | $-3$ | 2 | 0.64804 | Varied on a log scale $10^x$, $x$ is fit. $\phi_e = 4.447$ |
| $\kappa_{r,\tau}$ | molecules | 0 | 1000 | 536.2379 | |
| $\kappa_{\tau}$ | molecules | 0 | 1000 | 687.0312 | |
| $\kappa_{\gamma}$ | molecules | 0 | 1000 | 177.4952 | |
| $\kappa_H$ | molecules | 0 | $M_0/n_X$ | 15042.9702 | |
| $o_X$ | molecules | 0 | 1000 | 99.0688 | |
| $o_R$ | molecules | 0 | 1000 | 864.5958 | |
| $g_{T/E,0}$ | molecules | 10 | 100 | 14 | No. of promoters |
| $k$, $g_{H,0}$ scaling factor | unitless | 1 | 10 | 9.787 | $g_{H,0}$ is indirectly set $k \cdot g_{T/E,0} \therefore g_{H,0} = 132$ |
| $g_{P,0}$ | molecules | 0 | 100 | 2 | No. of promoters |
| $\phi_a$ | unitless | $-3$ | 2 | 0.98085 | Varied on a log scale $10^x$, $x$ is fit. $\phi_a = 9.57$ |
| $\phi_n$ | unitless | $-3$ | 2 | 1.2961 | Varied on a log scale $10^x$, $x$ is fit. $\phi_n = 19.77$ |

Table 2: **Known parameters**

| **Parameters** | **Value** | **Ref.** |
|---|---|---|
| $\phi_X$ | 10 | Calculated in [5] |
| $\phi_L$ | 6 | Calculated in [5] |
| $v_T$ | 726 molecules per min | Reported in [1] |
| $v_E$ | 5800 molecules per min | Reported in [1] |
| $k_{\{T,\ E\}}$ | 1000 molecules | Assumed |
| $g_{R,0}$ | 56 | Number of r-protein genes in *E. coli* |
| $g_{r,0}$ | 22 | Number of rRNA genes in *E. coli* |
| $q_X$ | 40 | Estimation based on data in [6] |
| $q_r$ | 50 | Estimation based on data in [6] |
| $n_{\{T,\ E,\ A,\ H\}}$ | 330 amino acids | Avg. *E. coli* gene is 1000 nucleotides [7] |
| $n_P$ | 3636 amino acids | Size of the *E. coli* RNA polymerase core complex |
| $n_R$ | 7459 amino acids | Size of the *E. coli* ribosome and associated factors |
| $n_r$ | 4566 nucleotides | Length of the in *E. coli* rRNA operon |
| $\beta_j$ | 1 cell/(molecule $\cdot$ min) | Assumed as in [1] |
| $\mu_j$ | 1 1/min | Assumed as in [1] |
| $b_j$ | 1 cell/(molecule $\cdot$ min) | Assumed as in [1] |
| $u_j$ | 1 1/min | Assumed as in [1] |
| $b_\rho$ | 1 cell/(molecule $\cdot$ min) | Assume diffusion limited as in [2] |
| $u_\rho$ | 1 1/min | Assume diffusion limited as in [2] |
| $\delta_{m_j}$ | 0.1 1/min | mRNA's have a half life of minutes [7] |
| $h_H$ | 4 | Assumed |
| $\tau_{r,max}$ | 5100 nucleotides/min | [6] |
| $\tau_{X,max}$ | 3300 nucleotides/min | [6] |
| $\gamma_{max}$ | 1260 amino acids/min | [6] |
| $M_0$ | $10^8$ amino acids | [7] |

RNA polymerase and ribosomes described in Equations 14 and 17. The definition of growth rate (Equation 25) is updated to include the additional translation complexes of the new genes. These modifications are carried out by expanding the set $j$ in the equations above to include the new genes.

We model the two gene RFP-GFP resource competition reporter circuit by introducing two new genes as described. Each gene, $i$, is regulated as follows:

$$\mathcal{R}_i = \frac{e \cdot u_i}{o_i + e} \tag{33}$$

where $u_i$ takes a value between 0 and 1 and scales the RNAP-promoter association rate.

##### 3.4 Simulating chemostat and batch culture

The different culture conditions can be simulated by selecting the form of Eq. 1 and 18. In the 'exponential growth' model, external nutrients are constant and we do not account for cell growth ($dN/dt = 0$). In the 'batch growth' model, growth rate is proportional to population size and the nutrient is depleted over time (in proportion to nutrient import into the $N$ cells of the population). This recapitulates logistic growth of a batch culture (see [1]). We assume that stationary phase cultures can be modelled by simulating the batch culture model with growth and energy consumption until $s_x = 0$ (at which point the the population ceases to increase as translation ceases). However, this clearly neglects any biological processes which might govern the transition to stationary phase. To simulate the fixed growth rate achievable in a chemostat, we set the value of $\delta$. At steady state $dN/dt = 0$, then $\lambda = \delta_N$.

We model these growth settings by changing the following 'environmental' parameters:

- The nutrient quality/efficiency, $\phi_e$.
- The initial external substrate, $s_{x,0}$.
- The nutrient influx parameter, $k_{in}$.
- The cellular decay and external substrate efflux parameter $\delta$.

In the exponential setting:

- $\phi_e$ is varied to mimic growth in different nutrient conditions (i.e. different 'qualities').
- $s_{x,0}$ is constant – $s_{x,0}$ remains constant as $ds_x/dt = 0$ in the exponential setting.
- $k_{in}$ and $\delta$ are unused in this model.

In the chemostat setting:

- $\phi_e$, $s_{x,0}$ and $k_{in}$ are set at constant values.
- $\delta$ is set to chosen growth rate, $\lambda_{set}$
